# Regulatory dynamic enzyme-cost flux balance analysis: A unifying framework for constraint-based modeling

**DOI:** 10.1101/802249

**Authors:** Lin Liu, Alexander Bockmayr

## Abstract

Integrated modeling of metabolism and gene regulation continues to be a major challenge in computational biology. While there exist approaches like regulatory flux balance analysis (rFBA), dynamic flux balance analysis (dFBA), resource balance analysis (RBA) or dynamic enzyme-cost flux balance analysis (deFBA) extending classical flux balance analysis (FBA) in various directions, there have been no constraint-based methods so far that allow predicting the dynamics of metabolism taking into account both macromolecule production costs and regulatory events.

In this paper, we introduce a new constraint-based modeling framework named *regulatory dynamic enzyme-cost flux balance analysis* (r-deFBA), which unifies dynamic modeling of metabolism, cellular resource allocation and transcriptional regulation in a hybrid discrete-continuous setting.

With r-deFBA, we can predict discrete regulatory states together with the continuous dynamics of reaction fluxes, external substrates, enzymes, and regulatory proteins needed to achieve a cellular objective such as maximizing biomass over a time interval. The dynamic optimization problem underlying r-deFBA can be reformulated as a mixed-integer linear optimization problem, for which there exist efficient solvers.

## 1 Introduction

Constraint-based modeling approaches have become a powerful tool to analyze genome-scale metabolic network reconstructions [1, 2]. Based on the steady-state assumption for internal metabolites, constraint-based methods like *flux balance analysis* (FBA) [3] use the stoichiometry of the metabolic network to define a feasible solution space of possible steady-state flux distributions. By choosing an objective function such as maximizing biomass, an optimal flux distribution can be predicted by solving a linear optimization problem (LP).

While standard FBA requires very few data, it is not able to capture more complex phenomena such as dynamics, resource allocation, or gene regulation. Extending work by Palsson et al. [4], Mahadevan et al. [5] in 2002 introduced *dynamic flux balance analysis* (dFBA) to maximize biomass production over a time interval, taking into account the dynamics of extracellular metabolites and biomass. To incorporate the synthesis costs of macromolecules, Goelzer et al. [6, 7] developed *resource balance analysis* (RBA), which allows predicting an optimal resource allocation for maximizing the steady-state growth rate. Independently, Lerman et al. [8, 9] introduced *ME-models*, a related approach for integrating metabolism and gene expression at steady-state. To combine these two ways of extending FBA, dynamics and resource allocation, several frameworks have been developed during the last years, which include *dynamic enzyme-cost FBA* (deFBA) [10], *conditional FBA* (cFBA) [11, 12], *dynamic resource balance analysis* (dRBA) [13] and *dynamicME* [14].

Concerning integrated modeling of metabolism and regulation, there exist approaches such as *regulatory flux balance analysis* (rFBA) [15] and FlexFlux [16], which combine Boolean or multi-valued logical rules for transcriptional regulation with a steady-state stoichiometric model of metabolism. Like the SOA variant of dynamic flux balance analysis [4, 5], these techniques iterate flux balance analysis by splitting the growth phase into discrete time steps. At each time step, the updated regulatory states are imposed as bounds on the reaction fluxes, while ignoring the costs for enzyme production. *Integrated FBA* (iFBA) [17] allows combining rFBA with a differential equation model for a specific subnetwork, while *integrated dynamic FBA* (idFBA) [18] brings together metabolism, regulation, and also signal transduction.

In addition to these iterative methods, there exist also static approaches for combining metabolism and gene regulation. *Steady-state regulatory flux balance analysis* (SR-FBA) [19] aims at studying the steady-state behaviors of a metabolic-regulatory network by adding Boolean rules to the linear constraints of FBA, resulting in a mixed-integer linear program [19]. *Probabilistic regulation of metabolism* (PROM) [20] makes use of microarray data sets to constrain the reaction upper bounds with a certain percentage of the maximal upper bound.

To summarize, we classify in Tab. 1 existing flux balance approaches according to whether or not they include dynamics, macromolecule production costs, and gene regulation. As can be seen from Tab. 1, there is currently no approach integrating all those features in a unifying framework.

**Table 1:** Constraint-based flux balance approaches

| Approaches | No regulation |  | Regulation included |  |
| --- | --- | --- | --- | --- |
|  | No enzyme costs | With enzyme costs | No enzyme costs | With enzyme costs |
| Static | FBA (1992) [21] | RBA (2011) [6]<br>ME models (2012) [8] | SR-FBA (2007) [19]<br>PROM (2010) [20] |  |
| Iterative | dFBA (SOA) (1994) [4, 5] | dynamicME (2019) [14] | rFBA (2001) [15]<br>iFBA (2008) [17] | idFBA (2008) [18] |
| Dynamic | dFBA (DOA) (2002) [5] | deFBA (2015) [10]<br>cFBA (2015) [11]<br>dRBA (2018) [13] |  | r-deFBA (this paper) |

In previous work [22], we introduced *metabolic-regulatory networks* (MRNs) to formalize the interplay of metabolism, macromolecule synthesis and gene regulation. To specify the dynamics of MRNs, we used a hybrid automata framework, combining continuous dynamics of metabolism with discrete control by regulatory events. In this formalization, the amounts of molecular species are represented by continuous variables. The discrete states of the system correspond to gene expression states of regulated proteins, which include regulatory proteins and regulated enzymes. In each discrete state, the continuous variables evolve according to a system of differential equations that is specific for this state. The guard conditions for the discrete state transitions depend on the amounts of the molecular species and associated thresholds.

In the present paper, we look at dynamic optimization or optimal control of the hybrid automata representing MRNs, which leads to a new constraint-based modeling framework called *regulatory dynamic enzyme-cost flux balance analysis* (r-deFBA). Like in other flux balance approaches, we apply a quasi steady-state assumption for the internal metabolites. The resulting dynamic optimization problem can be transformed into a mixed-integer linear optimization problem (MILP), for which there exist efficient solvers.

The organization of this paper is as follows: We start in Sect. 2 by recalling the definition of MRNs and the hybrid modeling framework from [22]. In Sect. 3, we formally introduce r-deFBA by formulating the metabolic constraints, the regulatory constraints, and the resulting dynamic optimization problem. To illustrate our approach, we consider two biological applications. In Sect. 4 we analyze the self-replicator model already considered in [22]. Finally, in Sect. 5, we apply our approach to a model of core carbon metabolism, inspired from [15] and [10].

## 2 Hybrid dynamics of metabolic-regulatory networks

Metabolic-regulatory networks (MRNs) were introduced in [22] to model in an integrated way metabolic reactions, transcriptional regulation, macromolecule production and structural components. In Fig. 1, we illustrate the schematic structure of a MRN model. By 𝒴 we denote the set of extracellular species with corresponding exchange fluxes **v**_*𝒴*_. The intermediate metabolites 𝒳 are transformed by intracellular fluxes **v**_*𝒳*_ and utilized to build macromolecules 𝒫 = 𝒬 ∪ ℰ ∪ ℛ 𝒫. For simplicity, we consider here only three types of macromolecules: non-catalytic compounds 𝒬 such as DNA and lipids, catalytic molecules ℰ including enzymes and ribosomes, and regulatory proteins ℛ 𝒫. The corresponding production fluxes are denoted by **v**_*ℰ*_, **v** _*𝒬*_ and **v**_*ℛ𝒫*_. Within ℰ = ℛℰ ∪ 𝒩 ℛℰ, we distinguish between regulated enzymes ℛℰ, and non-regulated enzymes 𝒩ℛℰ. Overall, the set of molecular species is defined as

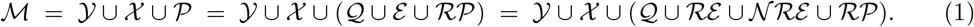

**Fig. 1:**
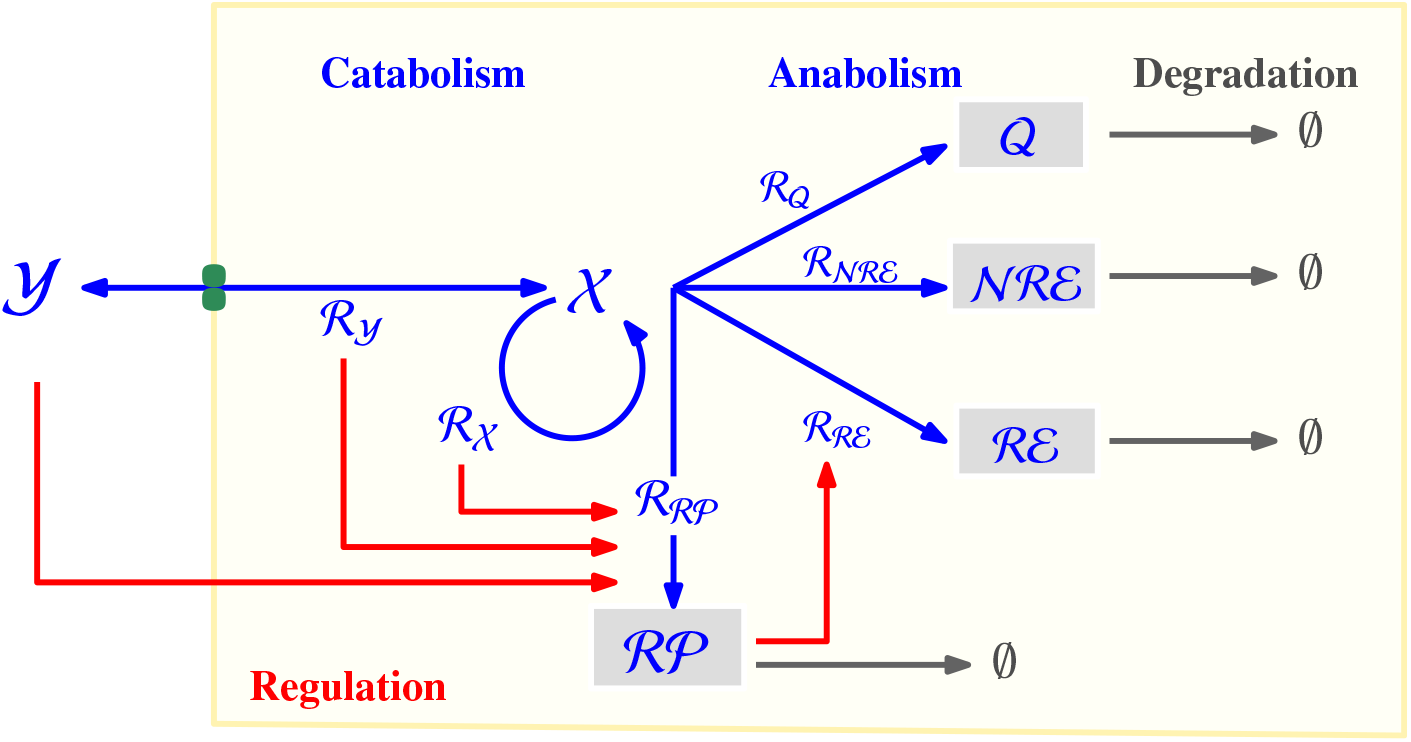
Schematic model of a metabolic-regulatory network (MRN). We distinguish three types of molecular species: extracellular species 𝒴, intermediate metabolites 𝒳, and macromolecules 𝒫, with corresponding vectors of molar amounts **Y, X, P**. By ℛ_*𝒴*_ we denote the set of exchange reactions with fluxes **v**_*𝒴*_. ℛ_*𝒳*_ is the set of intermediate reactions with the associated fluxes **v**_*𝒳*_. The macromolecules 𝒫 = 𝒬 ∪ ℛ ℰ ∪ 𝒩 ℛ ℰ ∪ ℛ 𝒫 are classified into quota compounds 𝒬, non-regulated enzymes 𝒩 ℛ ℰ, regulated enzymes ℛ ℰ, and regulatory proteins ℛ 𝒫, with vectors of molar amounts **Q, NRE, RE, RP**. Finally, ℛ _ℛ*ℰ*_, ℛ_*𝒩*ℛ*ℰ*_,ℛ_*𝒬*_,ℛ_ℛ*𝒫*_ and **v**_ℛ*ℰ*_, **v**_*𝒩*ℛ*ℰ*_, **v**_*𝒬*_, **v**_ℛ *𝒫*_ denote the sets of production reactions and corresponding fluxes.

### 2.1 Continuous dynamics

In a purely continuous modeling approach, the dynamics of the network would be described by a system of ordinary differential equations

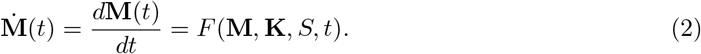

Following [10, 23, 22], we assume that **M**(*t*) denotes the molar amounts of the molecular species in ℳ at time *t*. Furthermore, **K** is a vector of kinetic parameters, *S* is the stoichiometric matrix, and *t* denotes time. The function *F* represents the kinetic laws that govern the dynamics, which could be mass action, Michaelis-Menten, Hill kinetics etc.

### 2.2 Discrete control

Continuous modeling of gene regulatory networks is known to be very difficult due to the lack of the necessary kinetic data. Therefore, we adopt a more qualitative approach to include regulation in our model. In this context, regulatory interactions, as illustrated by the red arrows in Fig. 1, refer to transcriptional regulation, i.e., we do not consider post-translational modifications. We assume that for each regulated protein *p* ∈ ℛ 𝒫 ∪ ℛℰ there are two possible states `on` and `off`, describing whether at a particular time *t* the gene encoding *p* is expressed or not.

Formally, we introduce a Boolean variable 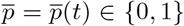 and a logical function *f*_*p*_ : ℝ^*n*^ → {0, 1}. Here, the Boolean value 0 corresponds to `off` and the value 1 to `on`. Each function *f*_*p*_ is defined as a Boolean combination (using the Boolean operations ¬ (not), ∧ (and), ∨ (or)) of atomic formulas of the form *x* ≥ *θ*, where *x* is a real variable and *θ* a threshold value. Overall, the regulation of our MRN is then formalized by a system of equations of the form

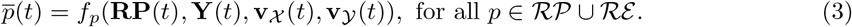

The logical function *f*_*p*_ indicates how the expression state of the regulated protein *p* depends on the current amounts of regulatory proteins, extracellular metabolites, and the reaction fluxes.

As an example, consider a regulatory rule stating that an enzyme *e* is activated by a regulatory protein *rp* above a certain threshold *θ* > 0. In this case, the expression state ē of *e* is given by the rule ē =*f*_*e*_ (*rp*(*t*)) = 1 if and only if *rp*(*t*) ≥ *θ*. Thus, whenever *rp*(*t*) ≥ *θ* holds, we have ē(t) = 1 (=`on`), while ē(t) = 0 (= `off`) whenever *rp*(*t*) < *θ*.

Note that in contrast to the metabolic-regulatory networks described in [22], the amounts of intermediate metabolites **X**(*t*) have been replaced in Eq. (3) with the flux values **v**_*𝒳*_ (*t*) and **v**_*𝒴*_ (*t*). This is due to the quasi steady-state assumption for internal metabolites, which is typical for constraint-based modeling approaches (see also [15]).

### 2.3 Hybrid system

Combining the continuous dynamics of metabolism in Eq. (2) with the discrete logical control in Eq. (3) leads to a hybrid discrete-continuous system, which we further explore in Sect. 3.

## 3 Formalization of r-deFBA

The r-deFBA framework that we propose in this paper aims at predicting from some initial conditions the continuous dynamics of metabolism and resource allocation together with discrete state transitions coming from genetic regulation. Compared with our earlier approach deFBA [10], regulatory logical constraints are included in addition to the metabolic constraints. Based on the schematic MRN model in Fig. 1 and the notation in Sect. 2, the metabolic and regulatory constraints of r-deFBA will now be described in detail.

### 3.1 Metabolic constraints

We start by recalling the constraints on metabolism, which are derived from dynamic enzyme-cost flux balance analysis (deFBA) [10, 23].

#### 3.1.1 Dynamics of external metabolites

The dynamics of the extracellular metabolites 𝒴 (nutrients and by-products) is modeled by a system of ordinary differential equations

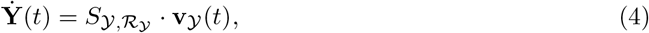

where 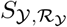 is a stoichiometric matrix in which the rows correspond to the extracellular metabolites 𝒴 and the columns to the exchange reactions ℛ_*𝒴*_. By **v**_*𝒴*_ (*t*) we denote the vector of exchange fluxes at time *t*.

#### 3.1.2 Dynamic production of macromolecules

The macromolecules 𝒫 = 𝒬 ∪ ℰ ∪ ℛ 𝒫 are assembled from metabolic precursors 𝒳 to ensure cellular survival and growth. The synthesis and the degradation of macromolecules is described by a system of differential equations

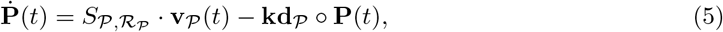

where 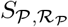 is a stoichiometric matrix in which the rows represent macromolecules and the columns macromolecule synthesis reactions. The vector **kd**_𝒫_ contains the degradation rates and ○ denotes the component-wise product of vectors.

#### 3.1.3 Quasi steady-state of intermediate metabolites

For the intermediate metabolites 𝒳 we assume that they are in quasi steady-state, i.e., the rate of production is equal to the rate of consumption. This leads to a system of algebraic equations

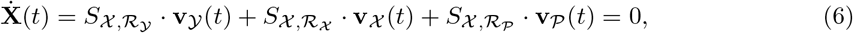

with stoichiometric matrices 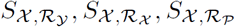 and fluxes **v**_*𝒴*_, **v**_*𝒳*_, **v**_*𝒫*_.

#### 3.1.4 Biomass composition constraint

In order to guarantee a sufficient production of non-catalytic macromolecules 𝒬 such as lipids and DNA, which are indispensable for cell growth and proliferation, we require that the mass of these quota compounds has to be at least a given fraction of the total biomass. Mathematically,

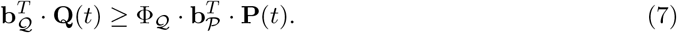

Here, **b**_*𝒫*_ is a vector with the molecular weights of all the macromolecules 𝒫, **b**_*𝒬*_ is the subvector of the molecular weights of the quota compounds 𝒬, the operation ·^*T*^ denotes transposition, and 0 < Φ_*𝒬*_ < 1 is a constant. ·^*T*^ denotes transposition, and 0 < Φ_**Q**_ < 1 is a constant.

#### 3.1.5 Enzymatic and translational capacity constraints

Fluxes through enzyme-catalyzed reactions are bound by the amount of the corresponding enzymes. If an enzyme catalyzes more than one reaction, the sum of all the reaction fluxes is limited by the enzyme amount. Formally, we get

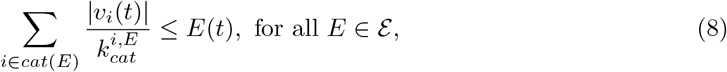

where *cat*(*E*) is the set of all reactions *i* catalyzed by enzyme *E* and 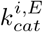 is the corresponding turnover rate. Note that this constraint also holds for protein translation and the ribosome, which is considered to be a special enzyme in our framework (cf. Sect. 2).

### 3.2 Regulatory logical control constraints

Extending the existing approaches for dynamic metabolic resource allocation such as deFBA, we now add two types of regulatory constraints. The first one describes the control of the discrete state transitions by the continuous variables, which corresponds to the triggering of the discrete jumps in the hybrid system. The second one is the control of the evolution of the continuous variables depending on the discrete state. Taken together, these two types of regulatory constraints specify the interplay between cellular regulation and metabolism.

#### 3.2.1 Control of discrete jumps

The key to the discrete dynamics of a hybrid system is how discrete state transitions are triggered by the continuous variables. According to Sect. 2, the expression state 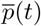 of all regulated proteins *p* is determined by a logical function *f*_*p*_ : ℝ^*n*^ → {0, 1} depending on the amounts of regulatory proteins, extracellular metabolites and reaction fluxes, i.e.,

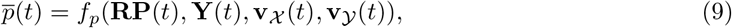

for all *p* ∈ ℛ𝒫 ∪ ℛℰ.

#### 3.2.2 Control of the continuous dynamics by the discrete states

While the regulatory constraints (9) describe how the discrete states depend on the continuous variables, we also have to specify how the continuous dynamics depends on the discrete state.

For a regulated protein *p*, the value of 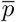 determines whether protein *p* is expressed or not. Therefore, if 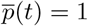, we impose as constraint that the production flux *v*_*p*_(*t*) should be at least *ε*_*p*_, while we require *v*_*p*_(*t*) to be zero in the case 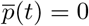. More formally, we get for all *p*∈ ℛ𝒫 ∪ ℛℰ the implications

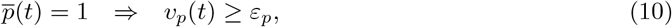

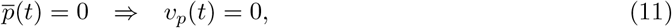

where *ε*_*p*_ > 0 is a lower bound for the production rate *v*_*p*_(*t*) of *p* in state `on`. We could also allow *ε*_*p*_ = 0 if we want to relax the model and determine the values of *v*_*p*_(*t*) by optimization like in deFBA.

Note that the values of the parameters *ε*_*p*_ significantly influence the dynamics of the system. Since the lower bounds constrain the production rates, they directly affect the abundances of the regulatory proteins and the regulated enzymes. Conversely, *p* is strictly constrained to be degraded whenever 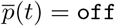.

### 3.3 Formulating r-deFBA as a dynamic optimization problem

Up to now, we have specified the metabolic constraints defining a dynamic solution space for cellular metabolism. In addition, we introduced the regulatory constraints to incorporate the dynamic interplay between gene regulation and metabolism. In order to predict how the cell can achieve optimal growth under these constraints, we formulate r-deFBA as a dynamic optimization problem, see Eq. 12. The objective is to compute time courses 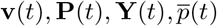 that maximize the total biomass production in a given time interval [*t*_0_, *t*_*f*_].

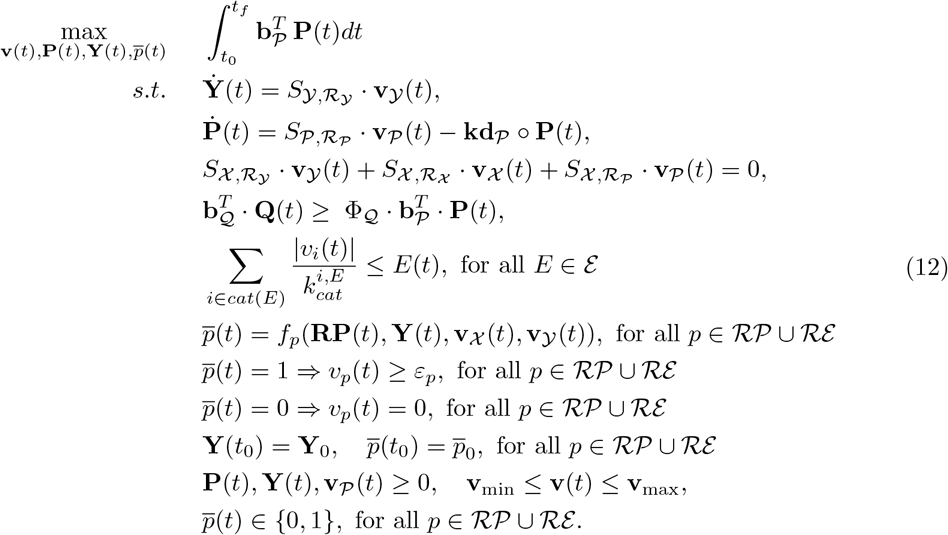

By **Y**_0_ and 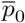 we denote the initial values of **Y**(*t*) and 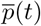 at time *t* = *t*_0_. In Eq. (12), the initial amounts **P**(*t*_0_) are variables whose values are determined by the dynamic optimization. Alternatively, initial values for **P**(*t*_0_) could also be precomputed using RBA.

Involving discrete and continuous variables, the r-deFBA problem in Eq. (12) can be reformulated as a mixed-integer linear optimization problem (MILP), for which there exist efficient solvers. To solve r-deFBA numerically, the dynamic real and Boolean variables are discretized in time like in [12]. The Boolean equations (9) and the logical implications (10)-(11) can be transformed into a system of linear inequalities using a standard recursive substitution procedure, see e.g. [19, 24]. For additional details we refer to the Appendix.

## 4 Case study 1: Regulatory self-replicator model

Carbon catabolite repression is a common phenomenon in bacteria, especially in *Escherichia coli* [25]. While these bacteria are able to grow on different carbon sources, they do not consume these in parallel, but one after the other. This is called diauxic growth and was described by Monod already in 1942. Mathematical modeling of diauxie has played an important role in understanding these phenomena [26].

As a possible model for diauxie, we built in [22] a small regulatory self-replicator network, extending earlier work in [15, 27]. To illustrate r-deFBA, we construct in Sect. 4.1 an r-deFBA model for this network. In Sect. 4.4 we compare the resulting dynamics for r-deFBA to standard deFBA and to the hybrid automata framework considered in [22].

### 4.1 Regulatory self-replicator network

The general workflow for building an r-deFBA model is illustrated in Fig. 2. Starting from a metabolic and a transcriptional regulatory network, we first construct a metabolic-regulatory network (MRN), as presented in Sect. 2.

**Fig. 2:**
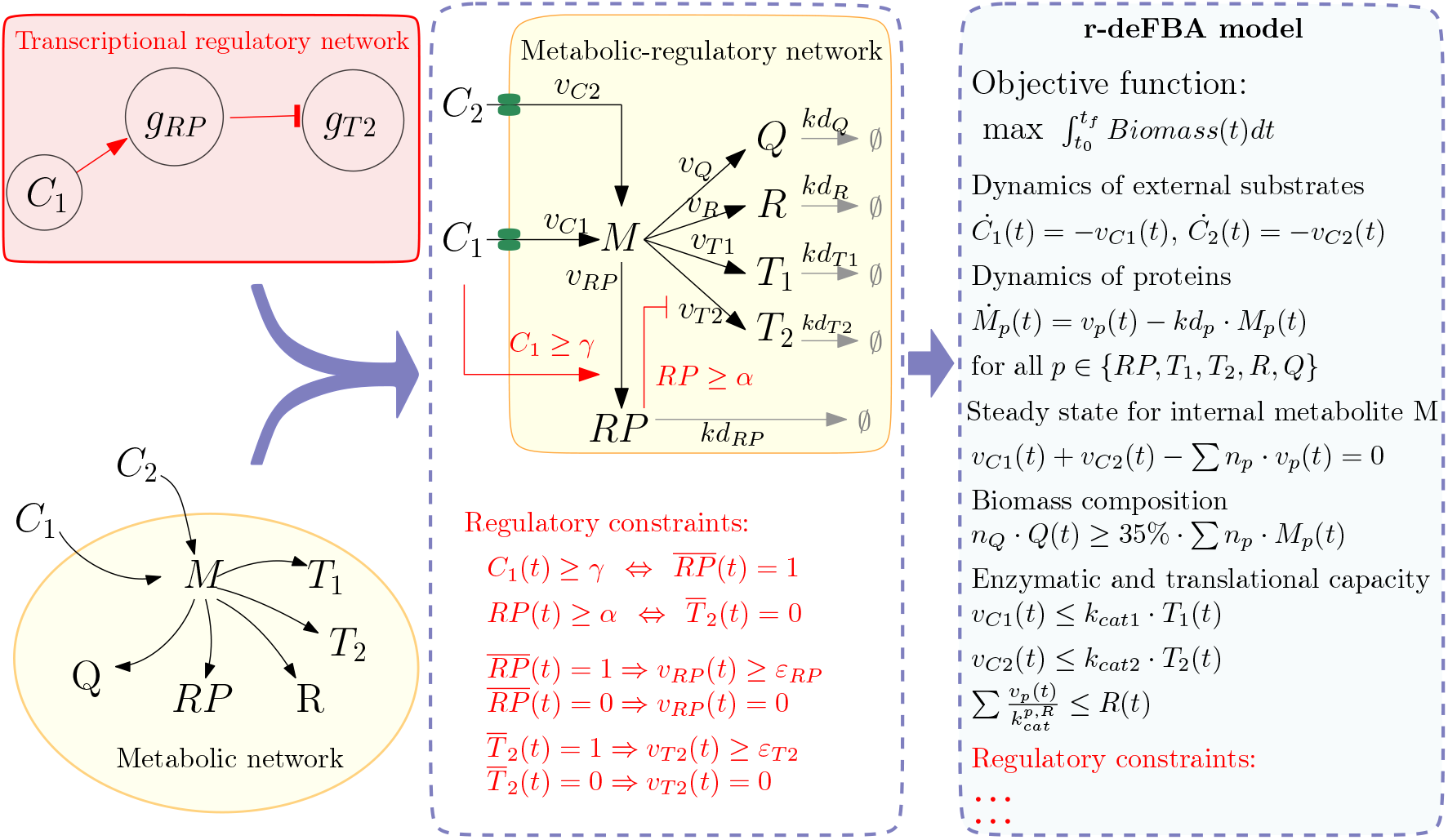
Workflow to build an r-deFBA model for the regulatory self-replicator with two regulatory rules. The Boolean variables 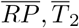 describe the expression state of the genes 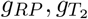, which determines the activity of the production reactions 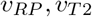. The thresholds 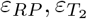 define the minimal expression levels for the regulated proteins *RP, T*_2_ to be in state `on`.

In the metabolic network of Fig. 2, we have two carbon sources 𝒴 = {*C*_1_, *C*_2_}, which are converted into precursor molecules 𝒳 = {*M*}. For simplicity, we assume only two uptake reactions *C*_1_ → *M, C*_2_ → *M*, catalyzed by enzymes *T*_1_ resp. *T*_2_. The precursor molecules *M* are used to synthesize five types of macromolecules 𝒫 = {*Q, R, T*_1_, *T*_2_, *RP*}, which are the enzymes *T*_1_, *T*_2_, regulatory proteins *RP*, housekeeping proteins *Q*, and ribosomes *R*. The stoichiometry of the synthesis reactions and corresponding parameter values are given in Tab. 2. The total biomass *Biomass*(*t*) is defined as the sum of the molecular masses

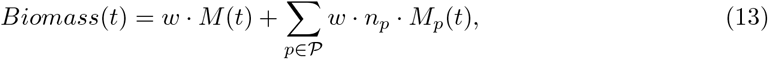

where *w* corresponds to the molar weight of one precursor molecule *M* and *n*_*p*_ is the number of precursor molecules needed to build one macromolecule *p*. By *M* (*t*) and *M*_*p*_(*t*) we denote again the molar amounts [mmol] of *M* resp. *p* ∈ 𝒫 at time *t*.

In the regulatory network of Fig. 2, *g*_*RP*_ and 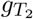 denote two genes encoding the regulated proteins *RP* and *T*_2_. We assume that *g*_*RP*_ is activated by the presence of *C*_1_ and that 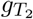 is inhibited by *g*_*RP*_. This leads to two regulatory rules

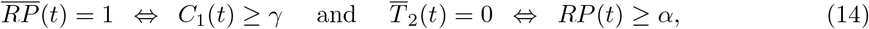

with thresholds *α, γ* > 0. The expression states 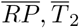 are linked to the flux variables *v*_*RP*_, *v*_*T* 2_ by the implications

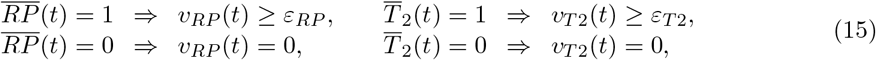

using thresholds *ε*_*RP*_, *ε*_*T* 2_ > 0.

**Table 2:** Metabolic and biomass reactions with corresponding parameters

| Metabolic reaction | Flux | Enzyme | Turnover rate [1/min] |  |
| --- | --- | --- | --- | --- |
| $C_1 \rightarrow M$ | $v_{C1}$ | $T_1$ | $k_{cat1} = 3000$ | |
| $C_2 \rightarrow M$ | $v_{C2}$ | $T_2$ | $k_{cat2} = 2000$ | |
| Biomass reaction | Flux | Enzyme | Turnover rate | Degradation rate |
| $n_p M \rightarrow p$ | $v_p$ | | $k_{cat}^{p,R} = k_r/n_p$ [1/min] | $kd_p$ [1/min] |
| $300M \rightarrow Q$ | $v_Q$ | R | 4.2 | 0.01 |
| $7459M \rightarrow R$ | $v_R$ | R | 0.1689 | 0.01 |
| $400M \rightarrow T_1$ | $v_{T1}$ | R | 3.15 | 0.01 |
| $1500M \rightarrow T_2$ | $v_{T2}$ | R | 0.84 | 0.01 |
| $300M \rightarrow RP$ | $v_{RP}$ | R | 4.2 | 0.2 |

### 4.2 r-deFBA and deFBA model

The full r-deFBA model of the regulatory self-replicator in Fig. 2 reads

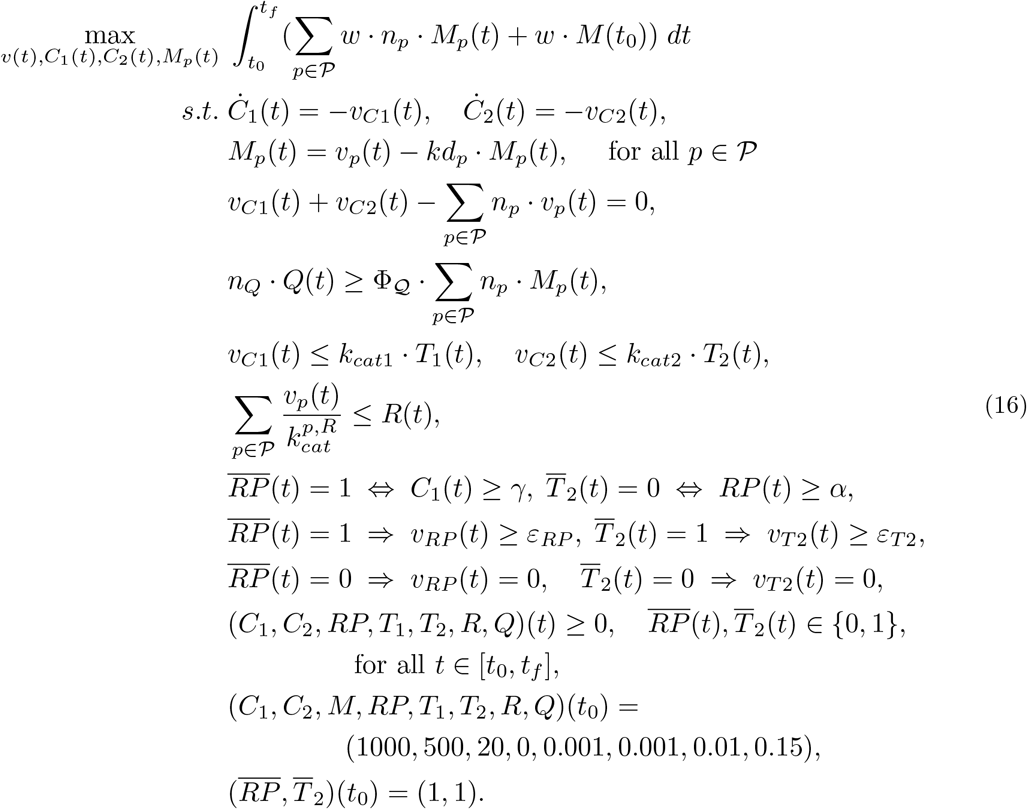

For the computations, we used the parameter values given in Tab. 2 and Tab. 3. The corresponding deFBA model is obtained by omitting the regulatory constraints.

**Table 3:** Additional parameters. Here, *k*_*r*_ denotes the elongation rate and *K*_*r*_, *K*_*T*_ are Michaelis constants.

| $t_0$ | $t_f$ | $w$ | $\Phi_Q$ | $\alpha$ | $\gamma$ | $\varepsilon_{RP}$ | $\varepsilon_{T2}$ | $K_T$ | $k_r$ | $K_r$ |
| --- | --- | --- | --- | --- | --- | --- | --- | --- | --- | --- |
| 0 | 55 | 100 | 0.35 | 0.03 | 20 | 0.01 | 0.01 | 1000 | 1260 | 7 |
| min | min | mg/mmol |  | mmol | mmol | mmol/min | mmol/min | mmol | 1/min |  |

### 4.3 Hybrid automaton

In [22], we also constructed a hybrid automaton for simulating the metabolic-regulatory network. The discrete states or locations correspond to the different 0-1 states of the Boolean variables 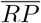 and 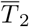, which means there are 2 × 2 = 4 discrete states. For the uptake of *C*_1_, *C*_2_ we assume a Michaelis-Menten kinetics

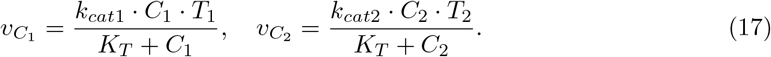

The parameter values can be found again in Tab. 2 and Tab. 3. Regarding the synthesis rate *v*_*p*_ of macromolecules *p* consisting of *n*_*p*_ precursors we assume a Michaelis-Menten type kinetics of the form

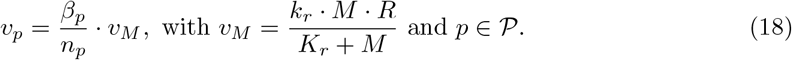

The weights *β*_*p*_ ≥ 0 denote the fraction of cellular resources allocated to protein *p*, with ∑*p β*_*p*_ = 1. For each location of the hybrid automaton, we assume for simplicity that the cellular resources are shared equally between the proteins that are expressed at this location. In other words, *β*_*p*_ = 1*/*3 in location 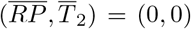, *β*_*p*_ = 1*/*4 in location (1, 0) resp. (0, 1), and *β*_*p*_ = 1*/*5 in location (1, 1). For further details, we refer to [22].

### 4.4 Comparing r-deFBA, deFBA, and the hybrid automaton

Next we compare the dynamics of the regulatory self-replicator obtained by r-deFBA, deFBA and the hybrid automaton, see Fig. 3. In all three simulations, we use the same parameter values, given in Tables 2 and 3, and the initial conditions from Eq. (16).

**Fig. 3:**
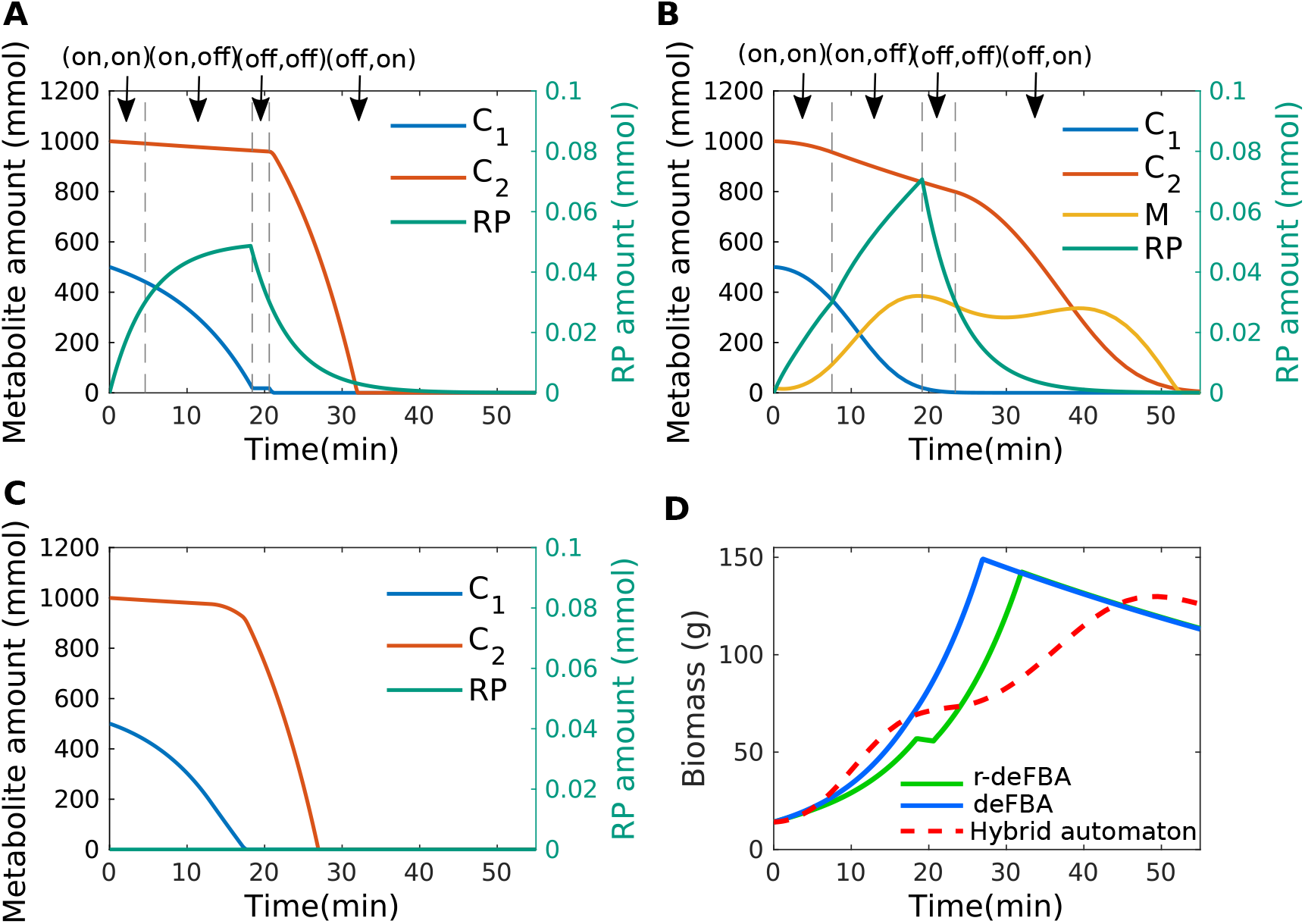
Time courses of *C*_1_, *C*_2_ (left axis) and *RP* (right axis) predicted by r-deFBA (**A**), the hybrid automaton (**B**), deFBA (**C**) and corresponding biomass production(**D**). In all three simulations, the same parameter and initial values were used. For r-deFBA and the hybrid automaton, we also indicate the discrete states 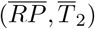, with the transitions marked by vertical dashed lines. Due to the quasi steady-state assumption, there is no trajectory for the precursor *M* in deFBA and r-deFBA.

The diauxic shift is predicted successfully by all three approaches. However, the underlying principles are different. By maximizing the biomass production while taking into account only the metabolic constraints, deFBA shows that the diauxic shift is an optimal metabolic behavior. In contrast, r-deFBA computes an optimal trajectory for biomass production, taking into account both the metabolic and the regulatory constraints. Due to the additional regulatory constraints, r-deFBA produces less biomass than deFBA and needs more time to consume the available carbon resources, see Fig. 3D. The continuous metabolic variables of the hybrid automaton evolve according to the Michaelis-Menten kinetics of Eq. (17) and (18). These kinetics depend on the current discrete state, which in turn is determined by the regulatory control, i.e., the jump conditions. As an optimal control strategy for the hybrid system representing the MRN, r-deFBA clearly gains more biomass than the hybrid automaton, but less than deFBA, which does not include regulation.

Both r-deFBA and the hybrid automaton successfully predict the discrete state transitions during diauxie. In Fig. 3A and 3B, the time profiles are divided into three growth phases, corresponding to the discrete state transitions. The transitions of r-deFBA are consistent with those obtained by the hybrid automaton. In the first growth phase, expression of *RP* is activated and 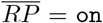 because initially *C*_1_ ≥ *γ*. We also have 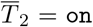 because *RP* is initialized by 0. Thus, the initial state of the network is 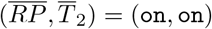. With time going on, *C*_1_ is consumed while *RP* is synthesized and accumulated. When *RP* reaches the threshold *α*, the synthesis of *T*_2_ is inhibited and the model jumps to the state (`on, off`). Next, when *C*_1_ < *γ*, the discrete state changes to (`off, off`), which represents the lag phase during diauxie. In this phase, enzyme *T*_2_ is still repressed until *RP* falls below its threshold *α*. Once this happens, the system switches to the final state (`off, on`), where *RP* < *α* and *T*_2_ is produced to metabolize *C*_2_. Overall, the interactions between discrete regulation and continuous metabolism are correctly incorporated in our r-deFBA. In deFBA, the regulatory protein *RP* remains at the initial value 0, see Fig. 3C. From the optimization perspective, there is no benefit in producing *RP* because it is a non-catalytic protein and does not sufficiently contribute to biomass.

Another interesting point is the production of macromolecules, see Fig. 4. In deFBA and r-deFBA, the dynamic optimization indicates that the production of *T*_1_ should be stopped once *C*_1_ is exhausted, although there is no regulatory control for *T*_1_. Intuitively, *T*_1_ is not needed anymore for uptake of *C*_1_. In order to increase biomass, it is better to produce *T*_2_ and *R*. When specifying the dynamics of the hybrid automaton in Eq. (18), we equally share the available resources between all synthesis reactions that are active in the current location. In real cells, this is unlikely to happen and not optimal for biomass production, as can be seen from Fig. 3D. Compared with the hybrid automaton, much more ribosome is produced in r-deFBA and deFBA, leading to a much larger biomass production, see Fig. 4. Clearly, how the cell allocates its resources to different enzymes will affect significantly the cellular growth.

**Fig. 4:**
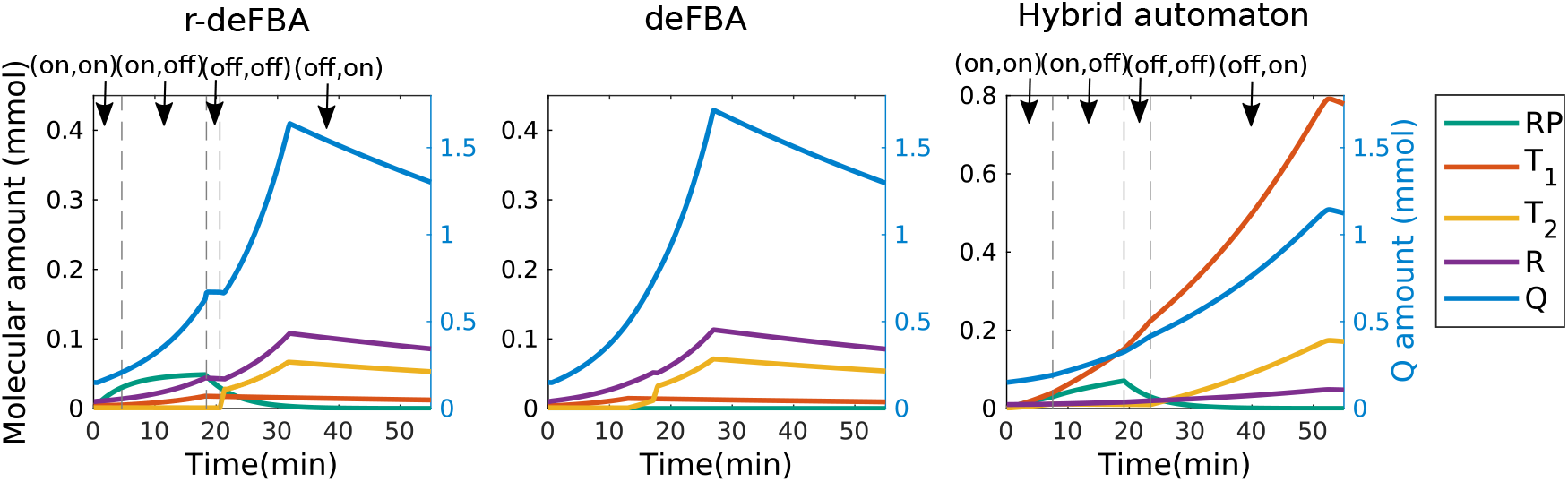
Time courses of *RP, T*_1_, *T*_2_, *R* (left axis) and *Q* (right axis) for r-deFBA, deFBA, and the hybrid automaton.

Since r-deFBA is more constrained than deFBA, the maximum biomass predicted by r-deFBA will always be less than or equal to the one for deFBA. However, the two maxima can get very close if the regulatory constraints are consistent with the objective in the dynamic optimization. In our simulation, deFBA successfully predicts the diauxic shift even without regulatory control, showing that this is an optimal strategy for biomass production. However, deFBA fails to provide information about how the cell should be regulated to achieve this result. In contrast, r-deFBA allows predicting both the dynamic evolution of regulatory proteins and the discrete state transitions which together enable the cell to implement an optimal growth strategy.

## 5 Case study 2: Core carbon metabolism

### 5.1 Core carbon metabolic network and related gene regulatory network

As before, we first construct a metabolic-regulatory network (MRN), see Fig. 5. Here we combine a metabolic and a regulatory network for core carbon metabolism based on [15, 10]. The metabolic network in Fig. 5A covers the major carbon pathways including glycolysis, TCA cycle, carbon storage, amino acid synthesis, pentose phosphate pathway, fermentation, and also the macromolecule synthesis.

**Fig. 5:**
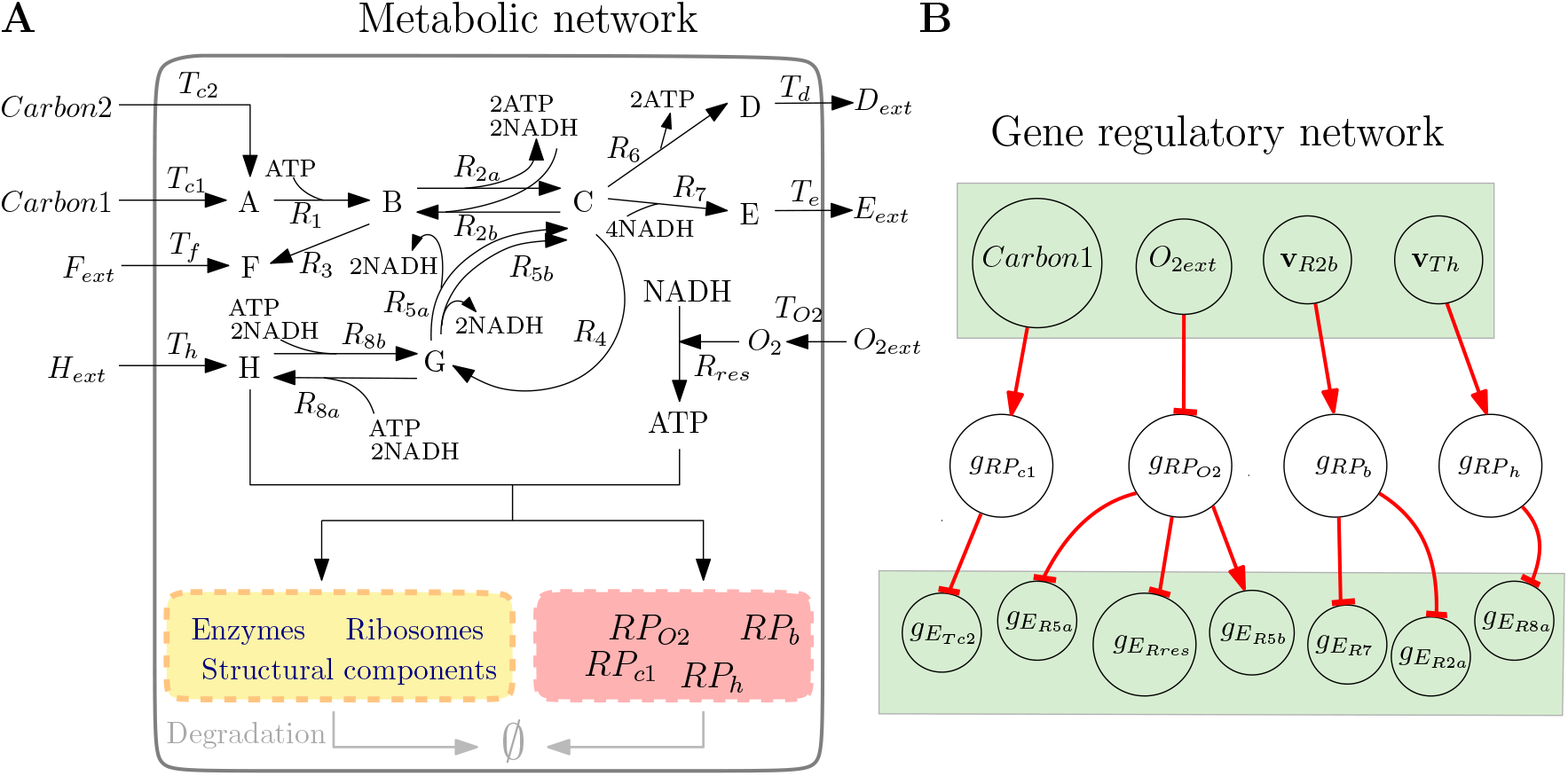
Metabolic network of core carbon metabolism (including macromolecule production) and corresponding gene regulatory network.

Using the notation from Sect. 2, we have the following molecular species:

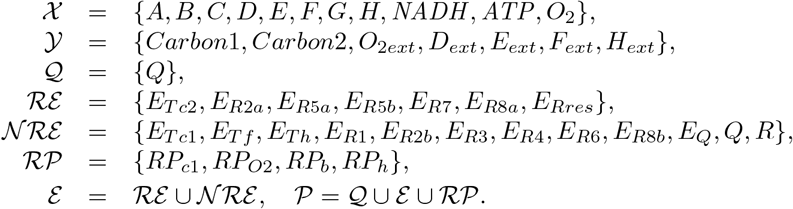

The details on the different metabolic and biomass reactions are given in Tab. 4 and 5. To get reasonable flux bounds on reactions describing diffusive exchange across the plasma membrane, we define the structural component *Q* as the enzymatic macromolecule for these reactions, together with an appropriate rate constant for diffusion [10].

**Table 4:** Metabolic reactions with catalyzing enzymes and turnover rates

| Reaction | Name | Enzyme | Turnover rate |
| --- | --- | --- | --- |
| | $i$ | $j$ | $k_{cat}^{i,j}$ [1/min] |
| Exchange reactions |  |  |  |
| $Carbon1 \rightarrow A$ | $T_{c1}$ | $E_{Tc1}$ | 3000 |
| $Carbon2 \rightarrow A$ | $T_{c2}$ | $E_{Tc2}$ | 2000 |
| $F_{ext} \rightarrow F$ | $T_f$ | $E_{Tf}$ | 3000 |
| $H_{ext} \rightarrow H$ | $T_h$ | $E_{Th}$ | 3000 |
| $O_{2ext} \rightarrow O_2$ | $T_{O2}$ | $Q$ | 1000 |
| $D \rightarrow D_{ext}$ | $T_d$ | $Q$ | 1000 |
| $E \rightarrow E_{ext}$ | $T_e$ | $Q$ | 1000 |
| Metabolic reactions |  |  |  |
| $A + ATP \rightarrow B$ | $R_1$ | $E_{R1}$ | 1800 |
| $B \rightarrow C + 2ATP + 2NADH$ | $R_{2a}$ | $E_{R2a}$ | 1800 |
| $C + 2ATP + 2NADH \rightarrow B$ | $R_{2b}$ | $E_{R2b}$ | 1800 |
| $C \rightarrow 2ATP + 3D$ | $R_6$ | $E_{R6}$ | 1800 |
| $C + 4NADH \rightarrow 3E$ | $R_7$ | $E_{R7}$ | 1800 |
| $B \rightarrow F$ | $R_3$ | $E_{R3}$ | 1800 |
| $C \rightarrow G$ | $R_4$ | $E_{R4}$ | 1800 |
| $G + ATP + 2NADH \rightarrow H$ | $R_{8a}$ | $E_{R8a}$ | 1800 |
| $H \rightarrow G + ATP + 2NADH$ | $R_{8b}$ | $E_{R8b}$ | 1800 |
| $G \rightarrow 0.8C + 2NADH$ | $R_{5a}$ | $E_{R5a}$ | 1800 |
| $G \rightarrow 0.8C + 2NADH$ | $R_{5b}$ | $E_{R5b}$ | 1800 |
| $O_2 + NADH \rightarrow ATP$ | $R_{res}$ | $E_{Rres}$ | 1800 |

Regarding the regulatory network of Fig. 5B, we identify again the state of a gene with the activity of the reaction producing the corresponding protein. For example, the gene state *g*_*RPc1*_ is identified with the activity state 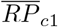 of the reaction producing *RP*_*c*1_. This means that the reaction synthesizing *RP*_*c*1_ will be active whenever the amount of external *Carbon*1 exceeds a given threshold. Conversely, the reaction will be blocked if not enough *Carbon*1 is available, see the regulatory rule for *RP*_*c*1_ in Tab. 5.

**Table 5:**
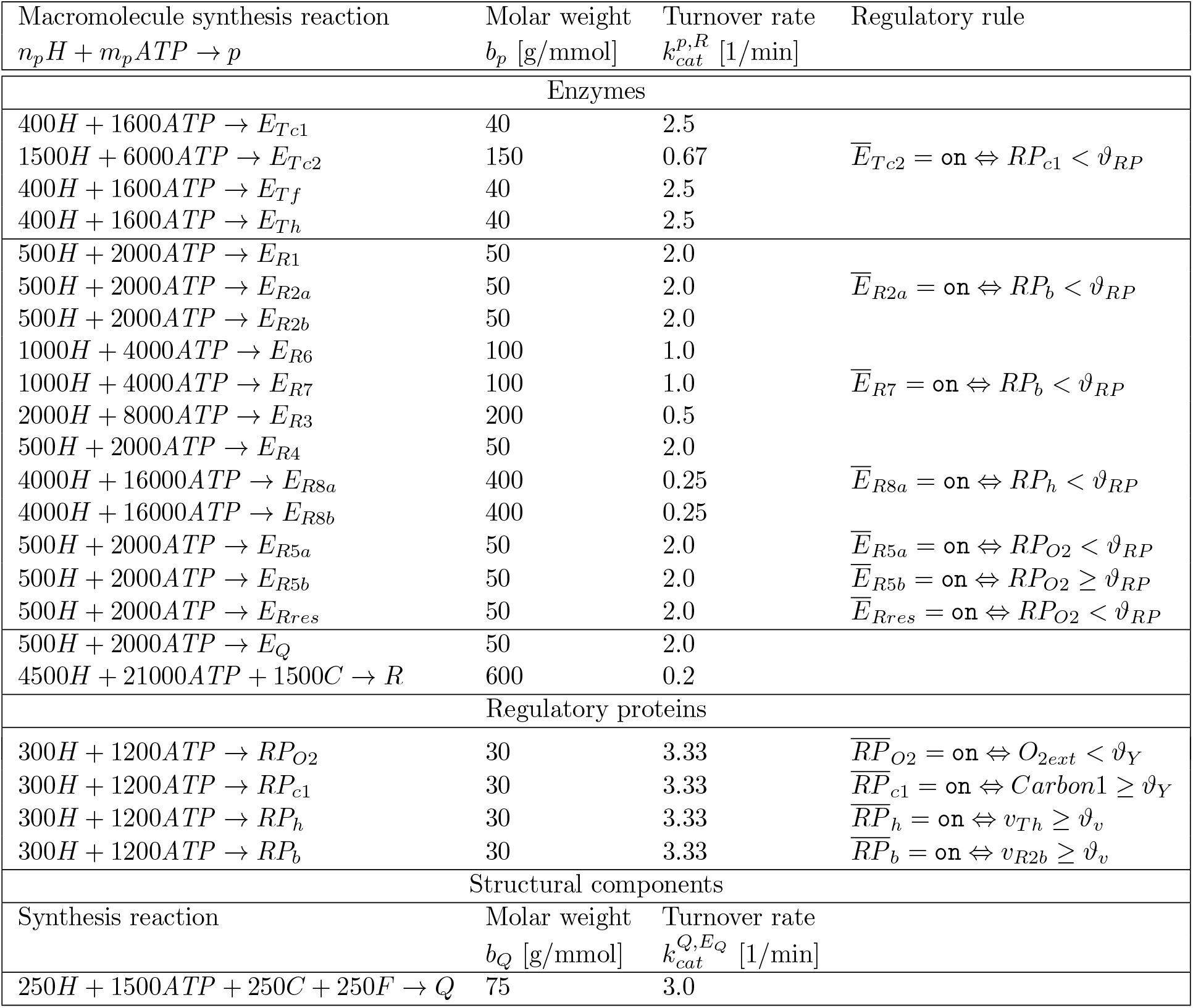
Macromolecule synthesis reactions with corresponding molar weights, turnover rates and regulatory rules

**Table 6:**
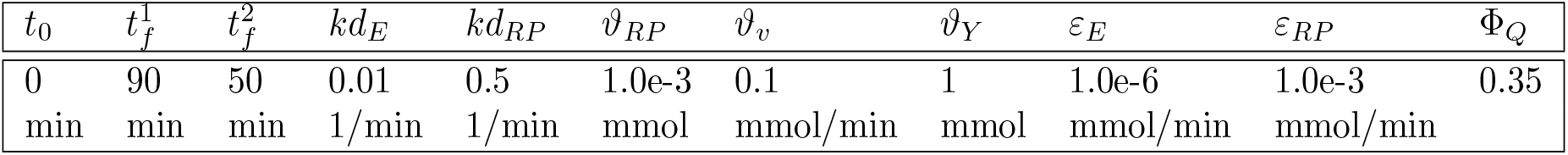
Parameters values for *E* ∈ *ε, RP* ∈ ℛ𝒫 and end times 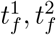 for Scenario 1 and 2.

Next we briefly describe the role of the four regulatory proteins *RP*_*c*1_, *RP*_*O*2_, *RP*_*h*_, *RP*_*b*_, for additional details we refer to [15]. The regulatory protein *RP*_*c*1_ is used to operate the diauxic shift between *Carbon*1 and *Carbon*2. Protein *RP*_*O*2_ is responsible for switching between the aerobic and anaerobic pathways, which are catalyzed by the isozymes *E*_*R*5*a*_, *E*_*R*5*b*_ respectively.

Protein *RP*_*h*_ regulates the usage of two primary nutrients of the cell, carbon sources and amino acid, via the reactions *R*_8*a*_ and *R*_8*b*_. These two reactions correspond to one reversible reaction that connects the TCA cycle with the uptake of extracellular amino acid *H*_*ext*_. Based on *RP*_*h*_, the cell will not produce amino acid from carbon sources by *R*_8*a*_ if enough amino acid can be taken up from the environment, i.e., if *v*_*Th*_ ≥ *ϑ*_*v*_.

The last regulatory protein *RP*_*b*_ is used to balance the key intermediate metabolites denoted by *B* and *C*. Since intracellular metabolite concentrations are not available due to the steady-state assumption, the reaction flux *v*_*R*2*b*_ is used as an internal signal. Production of *RP*_*b*_ is activated if *v*_*R*2*b*_ ≥ *ϑ*_*v*_, and *RP*_*b*_ abundance then inhibits the expression of *E*_*R*2*a*_ and *E*_*R*7_.

The complete r-deFBA model reads

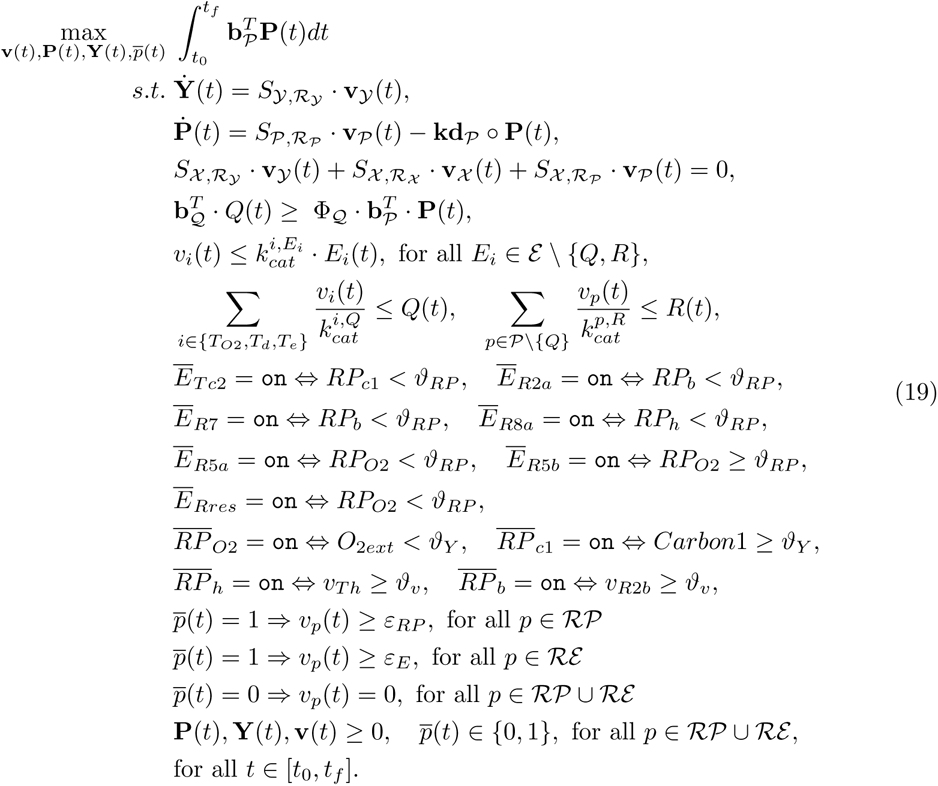

The initial values depend on the specific scenario and will be specified in the next section.

### 5.2 Comparing r-deFBA and deFBA

In total, there are 11 regulated proteins, which include 4 regulatory proteins and 7 regulated enzymes. The discrete state space thus contains 2<sup>11</sup> states, which are difficult to explore by the hybrid automaton. In the following, we present two scenarios to show how r-deFBA can be used to predict the integrated dynamics of metabolism and regulation even in a large state space. In each case, we compare r-deFBA with deFBA, which also models metabolism, but does not take into account the regulatory control.

#### 5.2.1 Scenario 1: Diauxie on two carbon sources

Our first scenario focuses again on the diauxie phenomenon. Initially, we set *Carbon*1 and *Carbon*2 to 1000 resp. 500 mmol, oxygen is given in excess, all other extracellular metabolites are set to 0. We do not specify the initial amounts of the macromolecules. Instead, these are computed by the optimization algorithm under the constraint that the initial biomass should be 1g. Thus the initial values for *t* = *t*_0_ are:

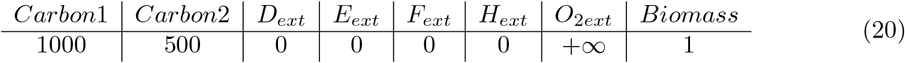

The Boolean variables are initialized by the 0-1 values for growth phase (a) in Tab. 7.

**Table 7:** Discrete state transitions in Scenario 1.

| Growth phase | $\overline{RP}_{c1}$ | $\overline{E}_{Tc2}$ | $\overline{RP}_{O2}$ | $\overline{E}_{R5a}$ | $\overline{E}_{R5b}$ | $\overline{E}_{Rres}$ | $\overline{RP}_b$ | $\overline{E}_{R2a}$ | $\overline{E}_{R7}$ | $\overline{RP}_h$ | $\overline{E}_{R8a}$ |
| --- | --- | --- | --- | --- | --- | --- | --- | --- | --- | --- | --- |
| a | 1 | 0 | 0 | 1 | 0 | 1 | 0 | 1 | 1 | 0 | 1 |
| b | 0 | 0 | 0 | 1 | 0 | 1 | 0 | 1 | 1 | 0 | 1 |
| c | 0 | 1 | 0 | 1 | 0 | 1 | 0 | 1 | 1 | 0 | 1 |

Comparing the results of r-deFBA and deFBA in Fig. 6A resp. 6C, we note that in both approaches *Carbon*1 is metabolized first. Yet, the biphasic increase of biomass is predicted only by r-deFBA, and not by deFBA. Although most of the available *Carbon*1 is utilized at the beginning, no lag phase is predicted by deFBA. The overall biomass production in the time interval [*t*_0_, *t*_*f*_] predicted by deFBA amounts to 111.3g, which is 8% more than the 103.0g obtained by r-deFBA.

**Fig. 6:**
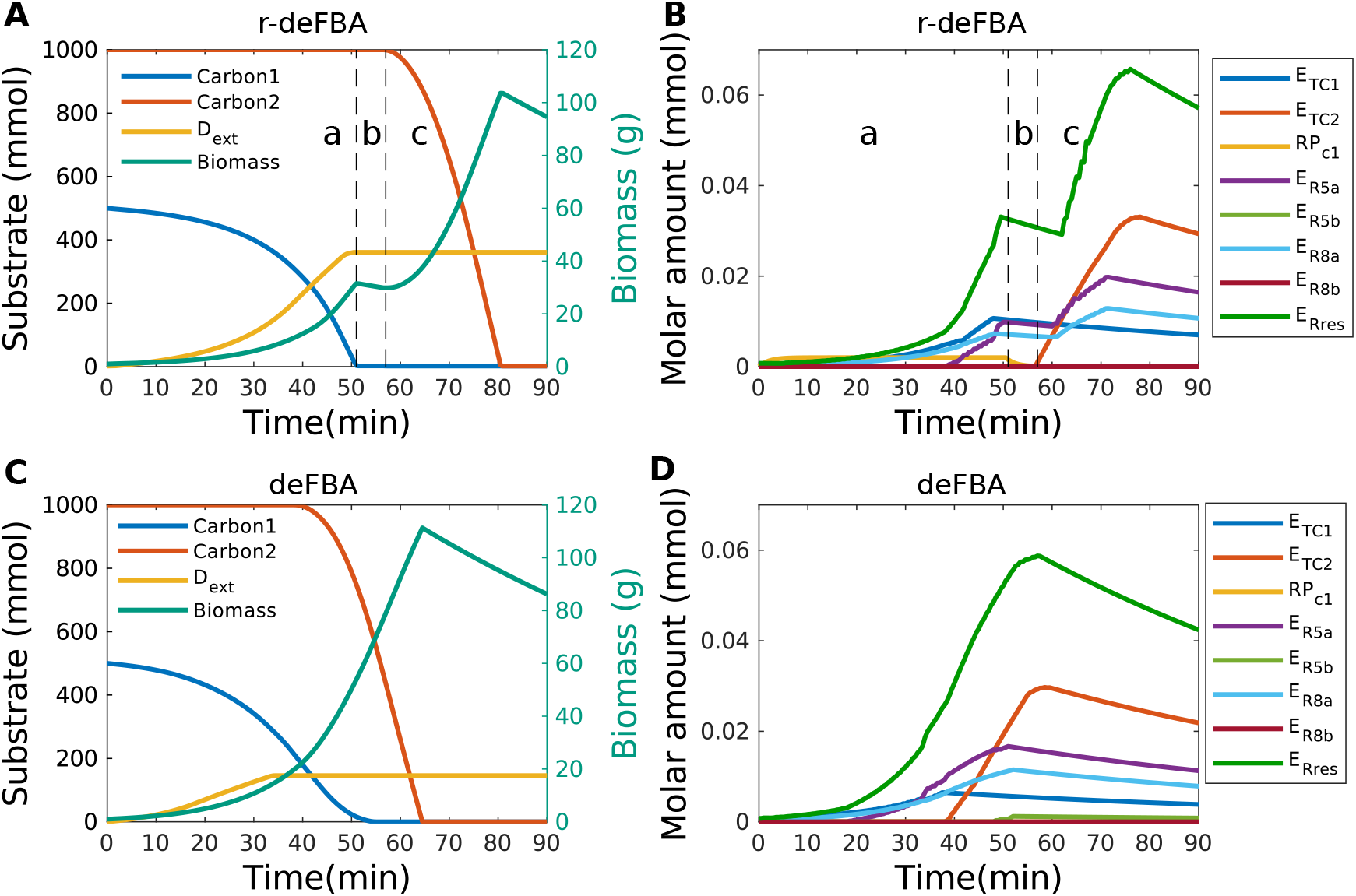
Dynamics of external substrates (left axis), total biomass (right axis) predicted by r-deFBA (**A**) and deFBA (**C**), and key regulated proteins (**B** and **D**) in Scenario 1.

At the level of individual proteins, 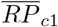 is produced in growth phase (a) of r-deFBA, for which *Carbon*1 ≥ *ϑ*_*Y*_, see Fig. 6B. Here, the expression of *Ē*_*Tc*2_ is inhibited since 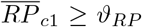. Thus, only *Carbon*1 supports growth during this period. Once it is exhausted, the growth shifts to phase (b). The indicator variable 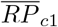 is triggered to be `off`, implying that *RP*_*c*1_ is degraded and not produced anymore. The expression of the transporter *E*_*Tc*2_ via *Ē*_*Tc*2_ is only activated when *RP*_*c*1_ < *ϑ*_ℛ𝒫_. So, no carbon can be taken up during phase (b) and the total biomass production shows a lag phase. Finally, in growth phase (c), the transporter *E*_*Tc*2_ is produced and biomass production resumes based on *Carbon*2.

Similarly in deFBA, *E*_*Tc*2_ is not synthesized as long as *Carbon*1 supports a high growth rate. The protein dynamics for r-deFBA and deFBA in Fig. 6B resp. 6D are also relatively close. However, *RP*_*c*1_ is not produced at all in deFBA and there is no lag phase, see Fig. 6D. In deFBA, the uptake of *Carbon*2 starts well before *Carbon*1 is exhausted, while in r-deFBA, *Carbon*1 and *Carbon*2 are metabolized one after the other, due to the regulatory control by *RP*_*c*1_. The synthesis of *RP*_*c*1_ generates extra costs in r-deFBA, such that the total biomass in r-deFBA is smaller than in deFBA.

The discrete state transitions for all the regulated proteins as predicted by r-deFBA are given in Tab. 7. Here we group together each of the four regulatory proteins with the corresponding regulated enzymes. The expression of the enzymes regulated by *RP*_*O*2_ does not change, since external oxygen is given in excess. Thus, 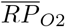 and *Ē*_*R*5*b*_ are always inhibited, while *Ē*_*R*5*a*_ and *Ē*_*Rres*_ remain activated. In Scenario 1, with no extracellular *H*_*ext*_ in the environment, reaction *R*_2*a*_ is constantly activated, consuming *Carbon*1, *Carbon*2, while *T*_*h*_ is inactive. Thus, the regulatory proteins 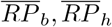 are always `off`, and *Ē*_*R*2*a*_, *Ē*_*R*7_, *Ē*_*R*8*a*_ are `on`. Note that the r-deFBA framework allows computing an optimal regulatory strategy for maximizing growth even though the discrete state space is very large.

The resource allocation during the carbon switch can also be investigated. In Fig. 7, we compare the dynamic biomass composition predicted by r-deFBA and deFBA for three kinds of macromolecules: structural components, enzymes and ribosomes. At the beginning, both approaches exhibit a stable biomass composition. The fraction of structural components initially stays around 35%, which corresponds to the lower bound imposed by the biomass constraint in Eq. (7). As *Carbon*1 is depleted, the structural components reach a rather high level, while the fractions of enzymes and ribosomes are decreasing in both predictions. Interestingly, in r-deFBA, the fractions of ribosomes and enzymes are increasing again while the structural components are going down at the outset of the second growth phase. This means that the cell has to allocate more resources to the ribosomes to start the second growth phase. In the last step, r-deFBA predicts a high fraction of structural components and a low fraction of enzymes and ribosomes, which can also be validated by experiments [28]. Overall, we obtain a biphasic resource allocation in r-deFBA, which is consistent with the two growth phases during diauxie. In deFBA, the quota fraction directly increases to about 80% of the total biomass and then keeps constant.

**Fig. 7:**
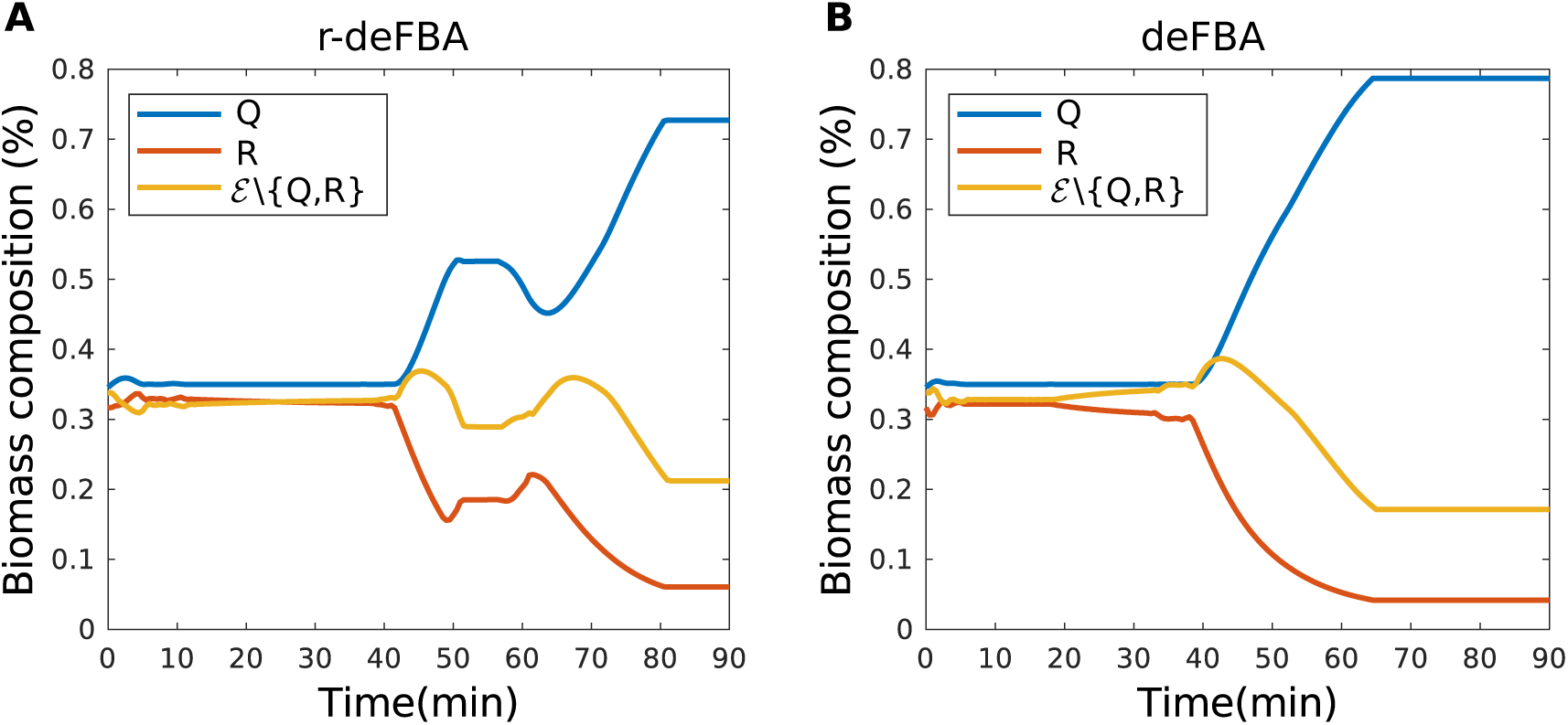
Dynamics of biomass composition in Scenario 1 with structural components *Q*, ribosomes *R* and enzymes *ε* \ {*Q, R*}.

#### 5.2.2 Scenario 2: growth on carbon and amino acid with amino acid in excess

Scenario 2 explores the dynamic growth on carbon and amino acid, with amino acid in excess. For *t* = *t*_0_ we choose the initial values:

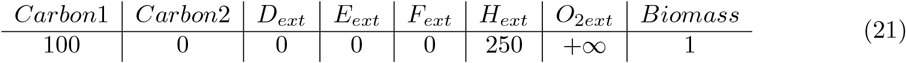

The Boolean variables are initialized by the 0-1 values for growth phase (a) in Tab. 8.

**Table 8:** Discrete state transitions in Scenario 2.

| Growth phase | $\overline{RP}_{c1}$ | $\overline{E}_{Tc2}$ | $\overline{RP}_{O2}$ | $\overline{E}_{R5a}$ | $\overline{E}_{R5b}$ | $\overline{E}_{Rres}$ | $\overline{RP}_b$ | $\overline{E}_{R2a}$ | $\overline{E}_{R7}$ | $\overline{RP}_h$ | $\overline{E}_{R8a}$ |
| --- | --- | --- | --- | --- | --- | --- | --- | --- | --- | --- | --- |
| a | 1 | 0 | 0 | 1 | 0 | 1 | 0 | 1 | 1 | 1 | 0 |
| b | 0 | 0 | 0 | 1 | 0 | 1 | 1 | 0 | 0 | 1 | 0 |
| c | 0 | 1 | 0 | 1 | 0 | 1 | 0 | 0 | 0 | 0 | 0 |

As can be seen from Fig. 8, both r-deFBA and deFBA first predict a co-utilization of *Carbon*1 and extracellular amino acid *H*_*ext*_, followed by the utilization of *H*_*ext*_ once *Carbon*1 has been exhausted. In the co-utilization phase, *R*_2*a*_ instead of *R*_2*b*_ is active to metabolize *Carbon*1. Consequently, enzyme *E*_*R*2*a*_ is synthesized in this phase, but not *E*_*R*2*b*_. When *Carbon*1 gets almost exhausted, enzyme *E*_*R*2*b*_ starts being produced in order to activate reaction *R*_2*b*_, see Fig. 8B and 8D. Now *B* has to be generated from *C*, since *B* is needed for growth. The switch between *R*_2*a*_ and *R*_2*b*_ is predicted by both approaches because it benefits growth. Although in deFBA the expression of the regulatory protein *RP*_*b*_ is not triggered to inhibit the synthesis of *E*_*R*2*a*_, the production of *E*_*R*2*a*_ stops in deFBA as well. Like *E*_*R*2*a*_, enzyme *E*_*R*8*b*_ also has a similar dynamics in both approaches.

**Fig. 8:**
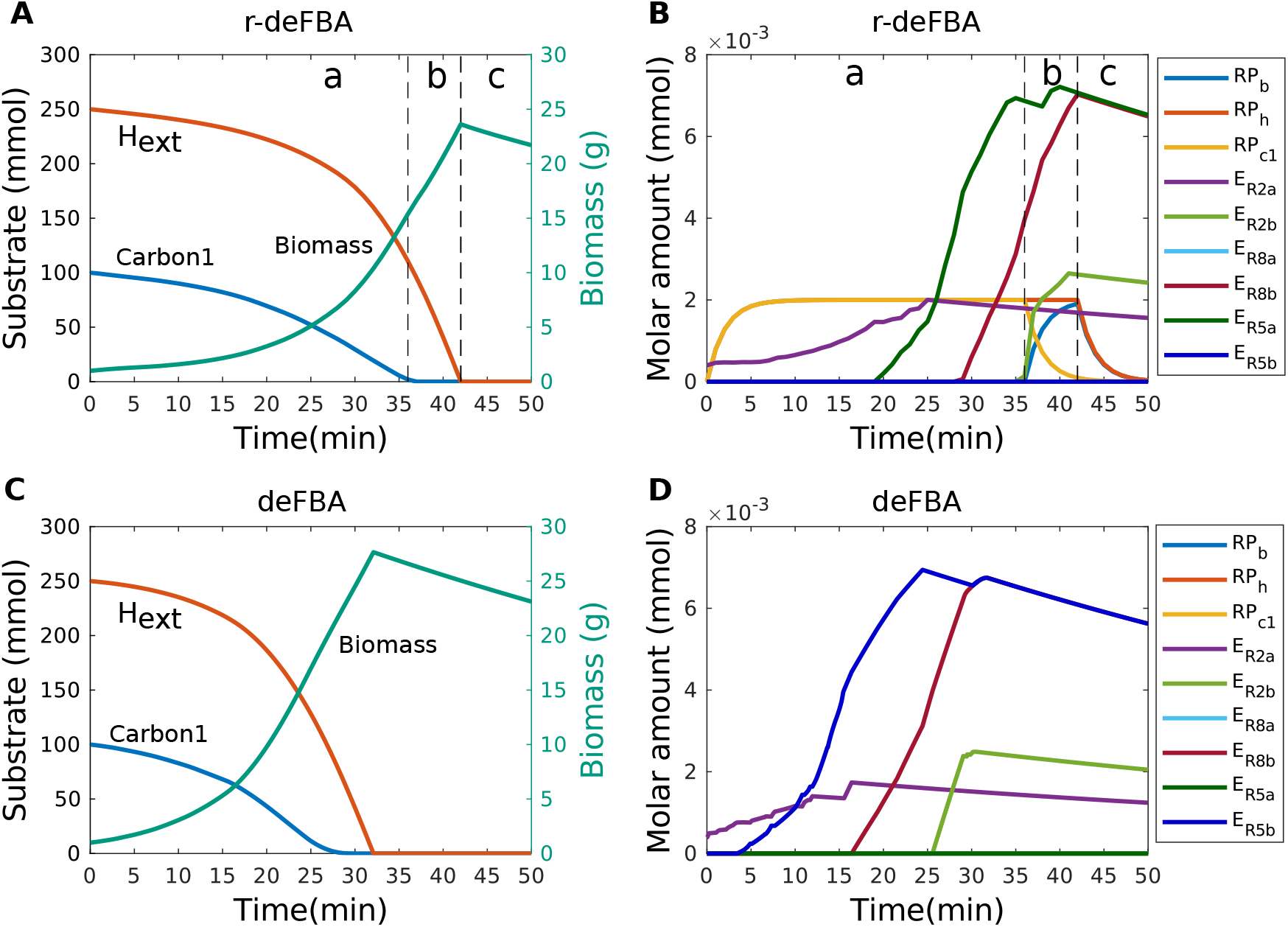
Dynamics of external metabolites (left axis) and biomass (right axis) predicted by r-deFBA(**A**) and deFBA (**C**), and key regulated proteins (**B** and **D**) in Scenario 2.

An interesting observation in the comparison is that deFBA activates enzyme *E*_*R*5*b*_ responsible for the anaerobic pathway, which is not consistent with the regulation. In contrast, r-deFBA produces enzyme *E*_*R*5*a*_ to catalyze reaction *R*_5*a*_, in agreement with the regulation by *RP*_*O*2_. Intuitively, *R*_5*a*_, *R*_5*b*_ are two alternative reactions that play the same role in the network, one in the aerobic, the other in the anaerobic case. Using only optimization without any regulatory information, deFBA cannot guarantee to choose the right pathway. Since optimal solutions are not unique, the solver can choose any of the two reactions or a combination thereof. Indeed, a small amount of *E*_*R*5*b*_ is produced by deFBA in the last phase of Scenario 1, even though this is not significant (see Fig. 6D). Clearly, the consistency between metabolism and regulation cannot be ensured by deFBA without additional regulatory information. In contrast, the dynamics of metabolism in r-deFBA highly depends on the activity of the regulatory proteins, and also the converse is true.

Switching between active reactions by r-deFBA is illustrated in Fig. 9. We can see in Fig. 9A that in the beginning *Carbon*1 and *H*_*ext*_ are co-utilized and *H* is obtained only from *H*_*ext*_. Since the model starts with a small biomass, even the TCA cycle is first inactive. Only when enzyme *E*_*R*5*a*_ has been synthesized, the TCA cycle is activated after 18 min, see Fig. 9B. Next in Fig. 9C, reaction *R*_8*b*_ is activated to furnish the TCA cycle with amino acid *H*, while releasing ATP and NADH. Now *Carbon*1 is not sufficient anymore to provide energy for growth. Finally, in Fig. 9D, *Carbon*1 has been exhausted, enzyme *E*_*R*2*b*_ is synthesized and *R*_2*b*_ is used to generate *B*.

**Fig. 9:**
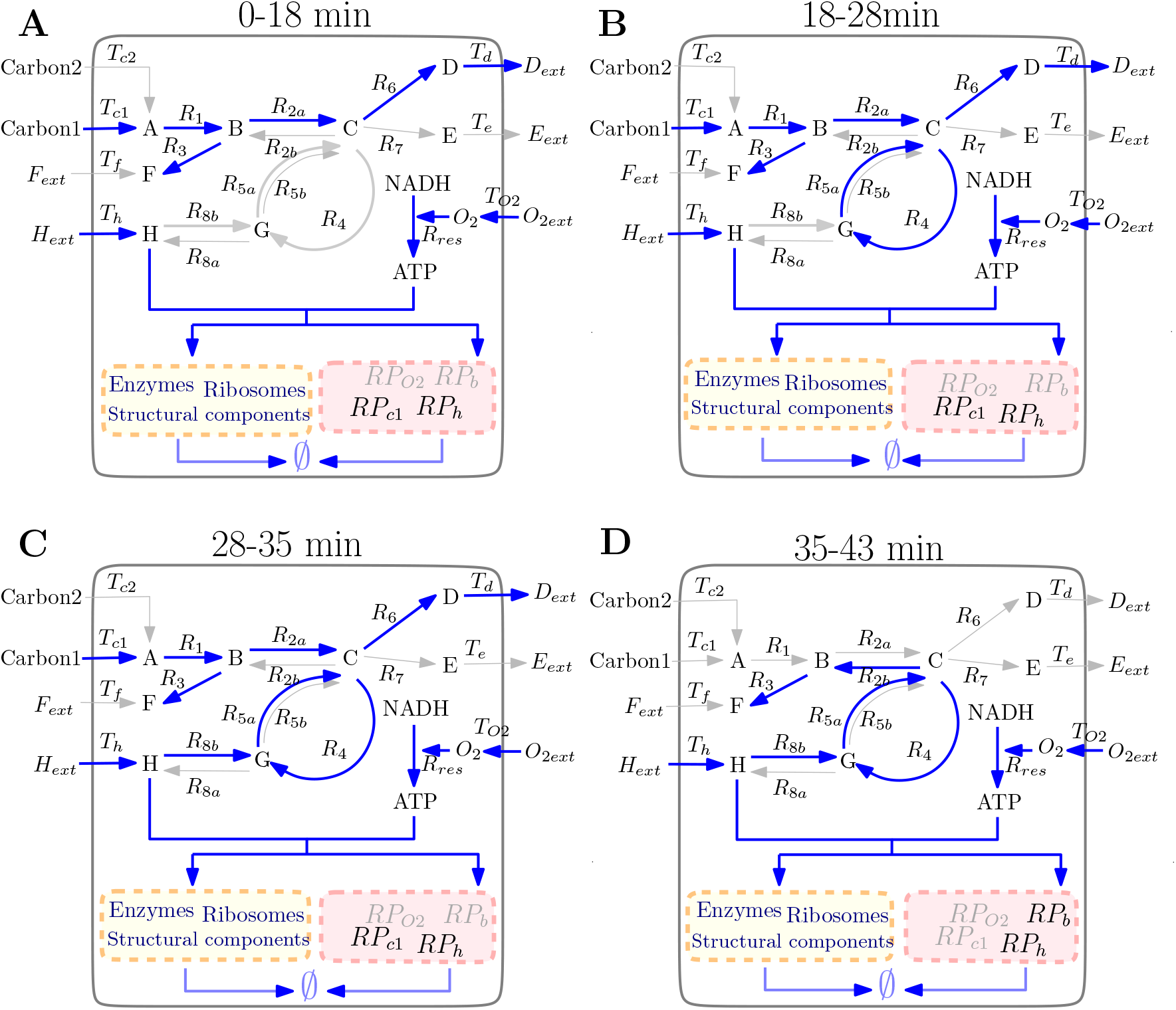
Patterns of active reactions predicted by r-deFBA in Scenario 2.

Regarding the discrete state transitions, we divide the simulation period of r-deFBA again into three phases (a), (b), and (c), see Fig. 8A and 8B. During the last phase (c), there is no growth since all the nutrients are exhausted. The key regulatory pathways analyzed in Scenario 2 are operated by the regulatory proteins *RP*_*b*_ and *RP*_*h*_. First, *R*_2*a*_ is active rather than *R*_2*b*_ for better metabolizing the carbon source. Hence, 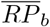 is `off` until *Carbon*1 is used up. Soon after growth phase (a), *R*_2*b*_ has to be activated to use *H*, so that *RP*_*b*_ is triggered to be produced (see Fig. 8B). The enzymes *Ē*_*R*2*a*_, *Ē*_*R*7_ then switch to `off`. Before *H*_*ext*_ has been exhausted, the expression state of *RP*_*h*_ is on because *T*_*h*_ has to be active for the uptake. Enzyme *Ē*_*R*8*a*_ is inhibited by *RP*_*h*_. The state transitions related with *RP*_*O*2_ are the same as in Scenario 1 because external oxygen is given in excess during the whole period. *Carbon*1 is given initially and exhausted at the end of growth phase (a). Although *Carbon*2 is set to 0 and the cell cannot use it, the expression state of *Ē*_*Tc*2_ is activated when *RP*_*c*1_ is totally degraded, due to the regulatory constraints. In Scenario 2, this happens by chance at the time when *H*_*ext*_ is used up. So, *E*_*Tc*2_ is on in phase (c). Meanwhile both *Carbon*1 and *H*_*ext*_ have been used up and the two reaction signals *v*_*Th*_ > *ϑ*_*v*_ and *v*_*R*2*b*_ > *ϑ*_*v*_ are inactive. The indicator variables 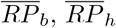 are turned `off` and the associated proteins *RP*_*b*_, *RP*_*h*_ are degraded in the last phase, see Fig. 8B.

## 6 Conclusion

Overall, r-deFBA computes optimal control strategies for hybrid automata representing metabolic-regulatory networks. Compared to previous approaches, in particular rFBA [15] and deFBA [10], r-deFBA allows for more realistic and accurate predictions by integrating the continuous dynamics of metabolism, including cellular resource allocation, with discrete regulatory control.

In purely discrete modeling frameworks for regulatory networks like the asynchronous logical formalism of R. Thomas [29], it is not possible to quantify the time delay between discrete state transitions. The hybrid automata approach proposed in [22] solves this problem by using continuous variables for regulatory protein amounts together with thresholds that trigger the discrete jumps. However, exploring the dynamics of these hybrid automata is difficult due to the exponentially large discrete state space. By computing an optimal control strategy for the hybrid automaton, r-deFBA is able to predict even in large state spaces the quantitative dynamics of the regulatory proteins together with the sequence of discrete state transitions that are needed to achieve optimal growth.

In summary, r-deFBA allows predicting optimal cellular resource allocation in a dynamic environment by integrating metabolic reactions, enzyme costs, quota compounds, and transcriptional regulation. Thus, r-deFBA considerably extends the predictive capabilities of current constraint-based modeling approaches as summarized in Tab. 1. Based on a hybrid discrete-continuous dynamics, r-deFBA is able to predict not only the continuous evolution of macromolecules and extracellular metabolites, but also the sequence of regulatory events needed to achieve an optimal growth. Finally, r-deFBA provides a solution for how to share enzymes between different reactions, which includes ribosome allocation in protein synthesis as a special case.

## Acknowledgements

The authors would like to thank M. Köbis, A. Reimers, and A. Röhl for fruitful discussions about this work. L. Liu gratefully acknowledges support from the China Scholarship Council (CSC) and DFG GRK 1772 “Computational Systems Biology”.

## A Appendix: Numerically solving r-deFBA as MILP

### Transforming logical constraints into linear inequalities

The dynamic optimization problem in (12) can be solved by transforming it into a mixed integer linear program (MILP). In a first step, the regulatory constraints (9)–(11) are converted into linear inequalities by standard reformulation techniques [30] that use additional 0-1 variables, which are called indicator variables. Note that the discrete states in the hybrid automata of MRNs are not affected by these indicator variables.

We introduce a vector of indicator variables **Ī**_ℛ𝒫_ ∈ {0, 1} ^ℛ𝒫^ (resp. **Ī**_𝒴_ ∈ {0, 1}^𝒴^) to indicate whether or not the regulatory protein amounts **RP** (resp. the extracellular metabolite amounts **Y**) are above the associated thresholds *θ*_ℛ𝒫_ (resp. *θ*_𝒴_). Mathematically

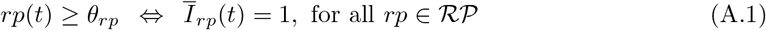

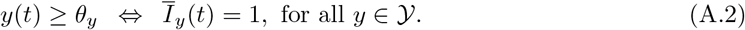

Eq. (A.1)–(A.2) can be transformed into linear inequalities by the reformulation

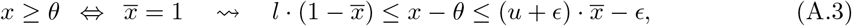

where *x* is a real variable, 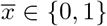 is the corresponding indicator variable, *θ* is the threshold, *l* resp. *u* is a lower resp. upper bound for *x* − *θ*, and *∊* is a small positive number.

By 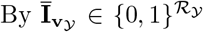 and 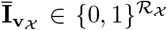 we describe the activity states of the reaction fluxes **v**_𝒴_ and **v**_𝒳_. A reaction is active if and only if the reaction flux is not zero, i.e., either strictly positive or stricly negative. Formally,

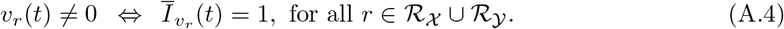

We transform these logical relations into linear inequalities by the reformulation

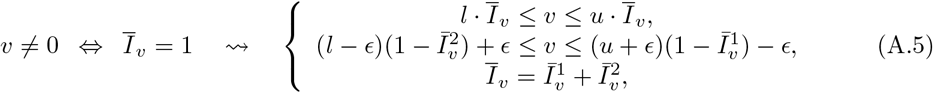

where *v* is a real variable, *Ī*_*v*_ ∈ {0, 1} is the corresponding indicator variable, *l* resp. *u* is a lower resp. upper bound for *v, ∊* is a small positive number, and 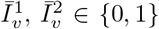 are two auxiliary 0-1 variables.

After introducing the indicator variables 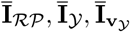 and 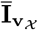, the regulatory constraints (9) are converted into Boolean equations

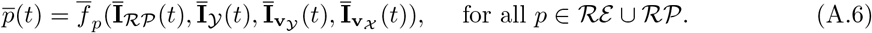

Similar to the original function *f*_*p*_ in (9), the Boolean function 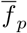 is defined in terms of the indicator variables 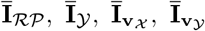 and the operations ¬ (not), ∧ (and), ∨ (or), such that 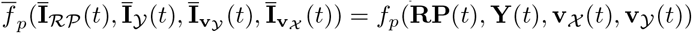.

Eq. (A.6) can be transformed into a set of linear inequalities by recursively applying the rules

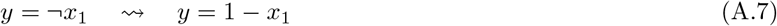

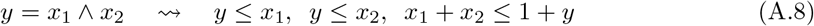

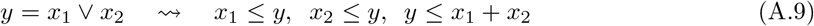

for variables *x*_1_, *x*_2_, *y* ∈ {0, 1} and by introducing additional 0-1 variables for the intermediate results.

Eq. (10)–(11) in the r-deFBA formulation are converted into linear inequalities by the reformulation

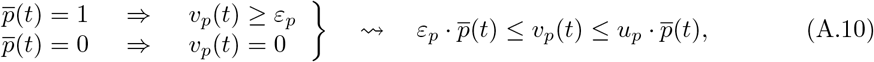

where *p* ∈ ℛ𝒫 ∪ℛ *ε* is a regulated protein, *ε*_*p*_ > 0 is lower bound on the flux *v*_*p*_(*t*) if *p* is expressed, and *u*_*p*_ is an upper bound on *v*_*p*_(*t*).

Using these transformations, the regulatory constraints can be reformulated as mixed 0-1 linear inequalities, see also [19, 24]. Together with the metabolic constraints, the dynamic r-deFBA optimization problem can be solved as a MILP by discretizing the continuous and Boolean variables in time.

### Discretizing the variables in time to solve r-deFBA

To solve numerically the dynamic optimization problem of r-deFBA, we discretize the variables in time, using the midpoint rule like in [12]. The macromolecular amounts **P** and extracellular metabolite amounts **Y** are discretized at each time point *t*_*k*_, *k* ∈ <u>{</u>0, …, *n*}. The flux variables **v** and the derivatives 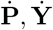 are discretized at the midpoint 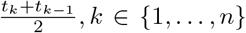, *k* ∈ {1, …, *n*}. In order to control the protein production, whose fluxes *v*_*p*_ are discretized at midpoint 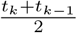, the Boolean variables 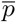, *p* ∈ ℛ𝒫 ∪ ℛ *ε*, are discretized at 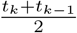, too. Indicator variables 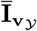 and 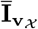 are also considered at the midpoint, which is consistent with the reaction fluxes **v**_𝒴_ and **v**_𝒳_. For **Ī**_𝒴_ and **Ī**_ℛ𝒫_, we calculate the corresponding amounts **Y** and **RP** at the time points *t*_*k*_ and *t*_*k*−1_ to determine the indicator values by

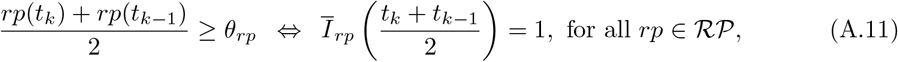

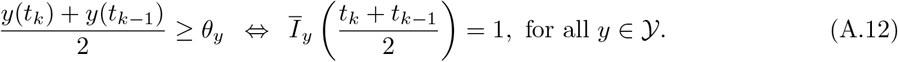

The resulting mixed-integer linear program (MILP) is given in Fig. 10. As we can see, all macromolecule amounts **P**, the derivatives 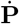 and all reaction fluxes **v** at each time point are continuous variables. The indicator variables of the extracellular species **Ī**_𝒴_, regulatory proteins **Ī**_ℛ𝒫_, reaction fluxes 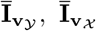 and expression states of regulated proteins 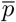 involved in the regulation are 0-1 variables. The biomass integral over the simulation period [*t*_0_, *t*_*f*_] is defined as the sum of all the macromolecule masses at each time point. Using this MILP, values for all the variables can be predicted using efficient MILP solvers such as Gurobi (http://www.gurobi.com), Cplex (https://www.ibm.com/products/ilog-cplex-optimization-studio) or SCIP (https://scip.zib.de/).

**Fig. 10:**
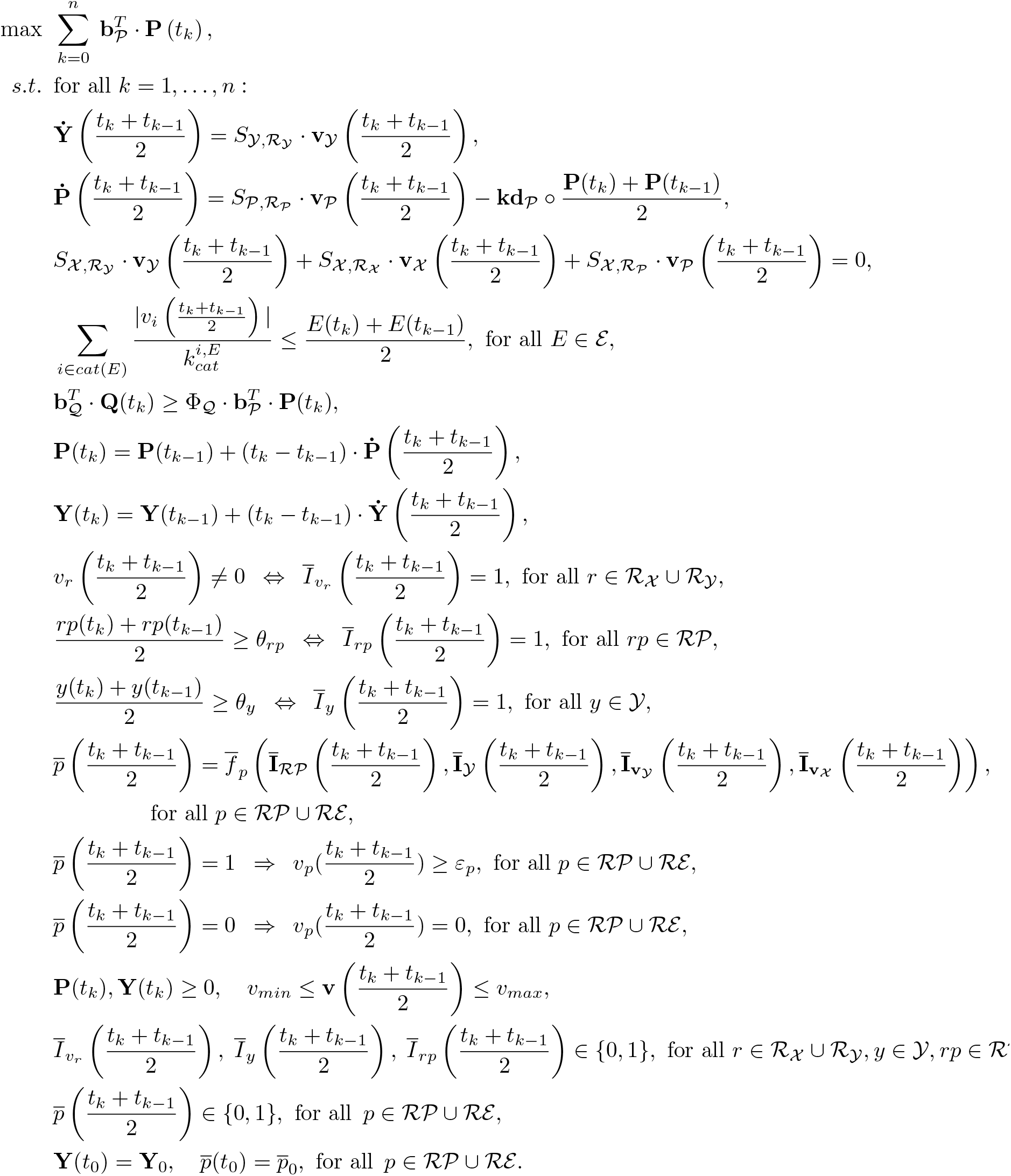
MILP for solving r-deFBA model

While the reformulation of r-deFBA as an MILP problem is our current solution strategy, developing possible alternative solution approaches is a topic of further research.

## References

[1] A. Bordbar, J. M. Monk, Z. A. King, and B. Ø. Palsson, “Constraint-based models predict metabolic and associated cellular functions,” Nat. Rev. Genet., vol. 15, pp. 107–120, Feb 2014.

[2] N. E. Lewis, H. Nagarajan, and B. Ø. Palsson, “Constraining the metabolic genotype-phenotype relationship using a phylogeny of in silico methods,” Nat. Rev. Microbiol., vol. 10, pp. 291–305, Feb 2012.

[3] J. D. Orth, I. Thiele, and B. Ø. Palsson, “What is flux balance analysis?,” Nat. Biotechnol., vol. 28, pp. 245–248, Mar 2010.

[4] A. Varma and B. Ø. Palsson, “Stoichiometric flux balance models quantitatively predict growth and metabolic by-product secretion in wild-type Escherichia coli W3110,” Appl. Environ. Microbiol., vol. 60, pp. 3724–3731, Oct 1994.

[5] R. Mahadevan, J. S. Edwards, and F. J. Doyle, “Dynamic flux balance analysis of diauxic growth in Escherichia coli,” Biophys. J., vol. 83, pp. 1331–1340, Sep 2002.

[6] A. Goelzer, V. Fromion, and G. Scorletti, “Cell design in bacteria as a convex optimization problem,” Automatica, vol. 47, no. 6, pp. 1210–1218, 2011.

[7] A. Goelzer, J. Muntel, V. Chubukov, M. Jules, E. Prestel, R. Nolker, M. Mariadassou, S. Aymerich, M. Hecker, P. Noirot, D. Becher, and V. Fromion, “Quantitative prediction of genome-wide resource allocation in bacteria,” Metab. Eng., vol. 32, pp. 232–243, Nov 2015.

[8] J. A. Lerman, D. R. Hyduke, H. Latif, V. A. Portnoy, N. E. Lewis, J. D. Orth, A. C. Schrimpe-Rutledge, R. D. Smith, J. N. Adkins, K. Zengler, and B. Ø. Palsson, “In silico method for modelling metabolism and gene product expression at genome scale,” Nat. Commun., vol. 3, p. 929, Jul 2012.

[9] E. J. O’Brien, J. A. Lerman, R. L. Chang, D. R. Hyduke, and B. Ø. Palsson, “Genome-scale models of metabolism and gene expression extend and refine growth phenotype prediction,” Mol. Syst. Biol., vol. 9, p. 693, Oct 2013.

[10] S. Waldherr, D. A. Oyarzun, and A. Bockmayr, “Dynamic optimization of metabolic networks coupled with gene expression,” J. Theor. Biol., vol. 365, pp. 469–485, Jan 2015.

[11] M. Rügen, A. Bockmayr, and R. Steuer, “Elucidating temporal resource allocation and diurnal dynamics in phototrophic metabolism using conditional FBA,” Sci Rep, vol. 5, p. 15247, 2015.

[12] A.-M. Reimers, H. Knoop, A. Bockmayr, and R. Steuer, “Cellular trade-offs and optimal resource allocation during cyanobacterial diurnal growth,” Proc. Natl. Acad. Sci. U.S.A., vol. 114, no. 31, pp. E6457–E6465, 2017.

[13] G. Jeanne, A. Goelzer, S. Tebbani, D. Dumur, and V. Fromion, “Dynamical resource allocation models for bioreactor optimization,” IFAC-PapersOnLine, vol. 51, no. 19, pp. 20–23, 2018.

[14] L. Yang, A. Ebrahim, C. J. Lloyd, M. A. Saunders, and B. Ø. Palsson, “DynamicME: dynamic simulation and refinement of integrated models of metabolism and protein expression,” BMC Syst Biol, vol. 13, p. 2, 01 2019.

[15] M. W. Covert, C. H. Schilling, and B. Ø. Palsson, “Regulation of gene expression in flux balance models of metabolism,” J. Theor. Biol., vol. 213, pp. 73–88, Nov 2001.

[16] L. Marmiesse, R. Peyraud, and L. Cottret, “FlexFlux: combining metabolic flux and regulatory network analyses,” BMC Syst Biol, vol. 9, p. 93, Dec 2015.

[17] M. W. Covert, N. Xiao, T. J. Chen, and J. R. Karr, “Integrating metabolic, transcriptional regulatory and signal transduction models in Escherichia coli,” Bioinformatics, vol. 24, no. 18, pp. 2044–2050, 2008.

[18] J. M. Lee, J. Min Lee, E. P. Gianchandani, J. A. Eddy, and J. A. Papin, “Dynamic analysis of integrated signaling, metabolic, and regulatory networks,” PLoS Comput. Biol., vol. 4, p. e1000086, May 2008.

[19] T. Shlomi, Y. Eisenberg, R. Sharan, and E. Ruppin, “A genome-scale computational study of the interplay between transcriptional regulation and metabolism,” Mol. Syst. Biol., vol. 3, no. 1, 2007.

[20] S. Chandrasekaran and N. D. Price, “Probabilistic integrative modeling of genome-scale metabolic and regulatory networks in Escherichia coli and Mycobacterium tuberculosis,” Proc. Natl. Acad. Sci. U.S.A., vol. 107, no. 47, pp. 17845–50, 2010.

[21] J. M. Savinell and B. Ø. Palsson, “Network analysis of intermediary metabolism using linear optimization. I. Development of mathematical formalism,” J. Theor. Biol., vol. 154, pp. 421–454, Feb 1992.

[22] L. Liu and A. Bockmayr, “Formalizing metabolic-regulatory networks by hybrid automata,” Acta Biotheor., July 2019.

[23] A.-M. Reimers, H. Lindhorst, and S. Waldherr, “A protocol for generating and exchanging (genome-scale) metabolic resource allocation models,” Metabolites, vol. 7, no. 3, p. 47, 2017.

[24] P. A. Jensen, K. A. Lutz, and J. A. Papin, “TIGER: Toolbox for integrating genome-scale metabolic models, expression data, and transcriptional regulatory networks,” BMC Syst Biol, vol. 5, no. 1, p. 147, 2011.

[25] B. Görke and J. Stülke, “Carbon catabolite repression in bacteria: many ways to make the most out of nutrients,” Nat. Rev. Microbiol., vol. 6, no. 8, p. 613, 2008.

[26] A. Kremling, J. Geiselmann, D. Ropers, and H. De Jong, “An ensemble of mathematical models showing diauxic growth behaviour,” BMC Syst Biol, vol. 12, no. 1, p. 82, 2018.

[27] D. Molenaar, R. Van Berlo, D. De Ridder, and B. Teusink, “Shifts in growth strategies reflect tradeoffs in cellular economics,” Mol. Syst. Biol., vol. 5, no. 1, 2009.

[28] E. Fischer and U. Sauer, “Large-scale in vivo flux analysis shows rigidity and suboptimal performance of Bacillus subtilis metabolism,” Nat. Genet., vol. 37, pp. 636–640, Jun 2005.

[29] R. Thomas and M. Kaufman, “Multistationarity, the basis of cell differentiation and memory. II. Logical analysis of regulatory networks in terms of feedback circuits,” Chaos: An Interdisciplinary Journal of Nonlinear Science, vol. 11, no. 1, pp. 180–195, 2001.

[30] H. P. Williams, Model building in mathematical programming-3rd ed. rev. Wiley, 1993.

